# Balancing Functional Tradeoffs between Protein Stability and ACE2 Binding in the SARS-CoV-2 Omicron BA.2, BA.2.75 and XBB Lineages : Dynamics-Based Network Models Reveal Epistatic Effects Modulating Compensatory Dynamic and Energetic Changes

**DOI:** 10.1101/2023.03.21.533701

**Authors:** Gennady Verkhivker, Mohammed Alshahrani, Grace Gupta

## Abstract

The evolutionary and functional studies suggested that the emergence of the Omicron variants can be determined by multiple fitness trade-offs including the immune escape, binding affinity for ACE2, conformational plasticity, protein stability and allosteric modulation. In this study, we systematically characterize conformational dynamics, protein stability and binding affinities of the SARS-CoV-2 Spike Omicron complexes with the host receptor ACE2 for BA.2, BA.2.75, XBB.1 and XBB.1.5 variants. We combined multiscale molecular simulations and dynamic analysis of allosteric interactions together with the ensemble-based mutational scanning of the protein residues and network modeling of epistatic interactions. This multifaceted computational study characterized molecular mechanisms and identified energetic hotspots that can mediate the predicted increased stability and the enhanced binding affinity of the BA.2.75 and XBB.1.5 complexes. The results suggested a mechanism driven by the stability hotspots and a spatially localized group of the Omicron binding affinity centers, while allowing for functionally beneficial neutral Omicron mutations in other binding interface positions. A network-based community model for the analysis of non-additive epistatic contributions in the Omicron complexes is proposed revealing the key role of the binding hotspots R498 and Y501 in mediating community-based epistatic couplings with other Omicron sites and allowing for compensatory dynamics and binding energetic changes. The results also showed that mutations in the convergent evolutionary hotspot F486 can modulate not only local interactions but also rewire the global network of local communities in this region allowing the F486P mutation to restore both the stability and binding affinity of the XBB.1.5 variant which may explain the growth advantages over the XBB.1 variant. The results of this study are consistent with a broad range of functional studies rationalizing functional roles of the Omicron mutation sites that form a coordinated network of hotspots enabling balance of multiple fitness tradeoffs and shaping up a complex functional landscape of virus transmissibility.

## Introduction

The staggering amount of structural and biochemical studies investigating mechanisms of SARS-CoV-2 infection have established a pivotal role of SARS-CoV-2 viral spike (S) glycoprotein in virus transmission and immune resistance [1–9]. The S protein consists of the intrinsically dynamic amino (N)-terminal S1 subunit that includes an N-terminal domain (NTD), the receptor-binding domain (RBD), and two structurally conserved subdomains SD1 and SD2 while the carboxyl (C)-terminal S2 subunit is structurally rigid. Stochastic conformational transformations between the closed (RBD-down) and open (RBD-up) forms of the S protein are orchestrated through coordinated global movements of the S1 subunit with respect to the largely immobilized S2 subunit, collectively eliciting diverse structural and functional adaptations of the S protein to various interacting partners, including binding with the host cell receptor ACE2 and immune responses to a wide spectrum of antibodies [10-15]. The thermodynamic and kinetic biophysical studies of the SARS-CoV-2 S functional trimers have revealed a complex interplay of local subdomain movements and long-range interactions that allow for subtle modulation of the RBD equilibrium and control of the relative populations of the open and closed states, thus regulating the exposure and binding affinities of the S protein with ACE2 and many interacting partners [16-18]. The cryo-EM and X-ray structures of the SARS-CoV-2 S variants of concern (VOC’s) in various functional states and complexes with antibodies revealed diversity of the binding epitopes and versatility of the S protein binding mechanisms with different classes of antibodies [19-28]. These studies unveiled that VOC mutations may act cooperatively to regulate a delicate balance and tradeoffs between various factors driving binding thermodynamics with ACE2 and immune evasion, while preserving the folding stability [29,30]. The biophysical thermostability studies of the D614G, BA.1, and BA.2 protein ectodomains demonstrated the reduced stability of the BA.1 RBD, while BA.2 RBD appeared to be more stable than BA.1 but less stable than the Wu-Hu-1 [31,32]. The cryo-EM structures of the S Omicron BA.1 trimers also suggested that in contrast to the original S strain with a mixture of open and closed conformations, the S Omicron BA.1 protein may adopt predominantly an open 1 RBD-up position predisposed for receptor binding [33-35]. In general, structural studies of the S Omicron BA.1 variant in complexes with ACE2 and various antibodies consistently indicated that evolutionary pressure may favor a mechanism in which the emerging mutations allow for an optimal balance between the enhanced ACE2 affinity and robust immune escape [36-41].

The recently reported structures of the BA.1.1, BA.2, and BA.3 RBD-ACE2 complexes pointed to a stronger binding of BA.1.1 and BA.2 subvariants as compared to BA.3 and BA.1 [42]. Structural and biochemical analysis of BA.2 binding with the human ACE2 (hACE2) showed that the S Omicron BA.2 trimer displayed binding affinity which was 11-fold higher than that of the S Wu-Hu-1 trimer and 2-fold higher than that of the S Omicron BA.1 [43]. Surface plasmon resonance (SPR) studies quantified the binding affinity of the Omicron BA.4/5 RBD for ACE2 which appeared to be stronger compared to the Wu-Hu-1 strain, BA.1, and BA.2 subvariants [44]. The cryo-EM structures and biochemical analysis of the S trimers for BA.1, BA.2, BA.3, and BA.4/BA.5 subvariants of Omicron reported the decreased binding affinity for the BA.4/BA.5 subvariants and the higher binding affinities for BA.2 as compared to other Omicron variants [45]. The thermal stability assays employed in this study also verified that S-trimers from BA.2 sublineages (B.2, B.2.12.1) were the least stable among B.1, B.2, B.3 and B.4 variants. Structure-functional studies of the Omicron BA.1, BA.2, BA.2.12.1, BA.4 and BA.5 subvariants showed the increased ACE2 binding affinity and stronger evasion of neutralizing antibody responses for BA.2 subvariants as compared to the Wu-Hu-1 and Delta strains, confirming that the compounded effect of the enhanced ACE2 receptor binding and stronger immune evasion may have contributed to the rapid spread of these Omicron sublineages [46].

A delicate balance between antibody evasion and ACE2 binding affinity was observed in biophysical studies of the Omicron BA.2.75 subvariant displaying a 9-fold enhancement of the binding affinity with ACE2 as compared to its parental BA.2 variant and showing the strongest ACE2 binding among all S variants measured to date [47]. This study discussed a possible role of N460K mutation in BA.2.75 in enhancing neutralization escape, while the reversed R493Q mutation featured in BA.2.75 subvariant could contribute to restoring the high ACE2 affinity but at the expense of the increased sensitivity to neutralization, overall suggesting that BA.2.75 mutations may prioritize ACE2 binding over immune escape. The cryo-EM conformations of the BA.2.75 S trimer in the open and closed forms as well as structures of the open BA.2.75 S trimer complexes with ACE2 reported thermal stabilities of the Omicron variants at neutral pH, showing that the BA.2.75 S-trimer was the most stable, followed by BA.1, BA.2.12.1, BA.5 and BA.2 variants [48]. The ACE2 binding affinities measured by SPR and reported in this study for Omicron subvariants BA.1, BA.2, BA.3, BA.4/5, BA.2.12.1, and BA.2.75 revealed that BA.2.75 displayed 4-6-fold increased binding affinity as compared to other Omicron variants. This study confirmed that BA.2.75 has the highest ACE2 affinity among all SARS-CoV-2 variants with the known experimental binding measurements [48]. Structure-functional investigation of BA.2.75 variant showed that NTD mutations K147E, F157L, and I210V, and two RBD substitutions N460K and R493Q can significantly increase infectivity, indicating that BA.2.75 variant can be endowed with significant antibody evasion potential while featuring the enhanced ACE2 binding as well as improved growth efficiency and intrinsic pathogenicity [49]. The receptor-binding assay performed in this study showed that G446S decreased the binding affinity of BA.2 RBD to ACE2 and was acquired to evade antiviral immunity while other substitutions in the BA.2.75 S RBD may compensate for this loss of binding affinity [49]. Similar balancing effects were observed in a study focusing on Omicron BA.4/5 [50] showing that the R493Q reversion in the BA.4/5 S protein could potentially contribute to evading immunity and marginal improvements in the ACE2 binding affinity while F486V substitution may have emerged to enforce immune evasion at the expense of the decreased ACE2 binding. Functional investigation of the BA.2.75 variant by examining mechanisms of virus infectivity and sensitivity to neutralizing antibodies revealed that N460K could be a key driver of the enhanced cell-cell fusion which enhances S processing, while G446S and N460K mutations may be responsible for the reduced neutralization sensitivity of BA.2.75 2 [51]. Although the mechanisms of infectivity may be different between BA.4/BA.5 and BA.2.75, these studies pointed to a unifying feature common to most mechanistic scenarios in which the acquisition of substitutions promoting immune evasion at the expense of the decreased ACE2 affinity is often counterbalanced by the emergence of mutations which compensate for this loss and promote the increased ACE2 binding [49-51].

The variants of past infection waves came from distinct branches of the SARS-CoV-2 family tree, but Omicron has spawned a series of subvariants that emerged from a single part of the SAR-CoV-2 evolutionary tree [52]. Among the emerging swarm (or soup) of latest SARS-CoV-2 variants, BQ.1.1 and XBB.1 variants have been circulating globally exhibiting superior growth advantages, where XBB.1.5 lineage particularly dominated with this subvariant making up to 28% of US COVID-19 cases, [52,53]. XBB.1 subvariant is a descendant of BA.2 and recombinant of BA.2.10.1 and BA.2.75 sublineages. XBB.1.5 is very similar to XBB.1 with a single RBD modification which is a notably rare two nucleotide substitution compared with the ancestral strain [52,53]. The biophysical studies of the S trimer binding with hACE2 for BA.2, BA.4/5, BQ.1, BQ.1.1, XBB, and XBB.1, variants showed that the binding affinities of BQ.1 and BQ.1.1 were comparable to that of BA.4/5 spike, while binding XBB and XBB.1 was similar to that of BA.2 variant [54]. According to this study, a moderate attenuation of the ACE2 binding affinity for XBB and XBB.1 variants could be attributed to F486S mutation removing the favorable interactions and inducing the decreased binding, and a compensatory R493Q mutation that partly restores the loss in the ACE2 binding [54]. It was also found that BA.2.75.2, BQ.1.1 and XBB.1 variants exhibited the lowest vaccine-elicited neutralization, thus indicating that emerging variants may have evolved to elicit stronger immune evasion without sacrificing ACE2 binding [54].

XBB.1.5, which is a subvariant of the recombinant mutant XBB, has shown a substantial growth advantage compared to both BQ.1.1 and XBB.1 [55-57]. The biochemical studies examined the binding affinity of the XBB.1.5 RBD to hACE2 revealing the dissociation constant K_D_ = 3.4 nM which was similar to that of BA.2.75 (K_D_=1.8 nM) while significantly stronger than that of XBB.1 (K_D_=19 nM) and BQ.1.1 (K_D_=8.1 nM) [56]. According to this study, XBB.1.5 is equally immune evasive as XBB.1 but may have growth advantage by virtue of the higher ACE2 binding affinity owing to a single S486P mutation as F486S substitution in XBB.1 can significantly compromise the local hydrophobic packing while F486P in XBB.1.5 subvariant can restore most of the favorable hydrophobic contacts [56]. Subsequent functional studies confirmed that the growth advantage and the increased transmissibility of the XBB.1.5 lineage may be a consequence of the retained neutralization resistance and the improved ACE2 binding affinity [57] These findings were consistent with the original deep mutational scanning (DMS) of the RBD residues using B.1, BA.1 and BA.2 backgrounds showing that F486 substitutions generally reduce ACE2 binding affinity due to decreased hydrophobic contacts, but these changes are more detrimental for F486S as compared to a modest loss for F486P [58,59]. The recent experimental studies examining the efficacy of COVID-19 vaccines and antibodies against XBB.1.5 subvariant disclosed that the neutralizing activity against XBB.1.5 was considerably lower than that against the ancestral strain and BA.2, while similar immune evasion potential was observed for XBB.1 and XBB.1.5 [60,61]. These studies confirmed that the high transmissibility and rapid surge of XBB.1.5 variant may be primarily due to the strong ACE2 binding affinity which is comparable only to the BA.2.75 variant, while retaining immune evasion similar to XBB.1 variant yields the overall better fitness tradeoff and leads to the observe growth advantages. The newly emerging variants such as BA.2.3.20, BA.2.75.2, BQ.1.1 and especially XBB, a recombinant of BJ.1 and BM.1.1.1 display substantial growth advantages over previous Omicron variants, and some RBD residues including R346, K356, K444, V445, G446, N450, L452, N460, F486, F490, R493 and S494 were found mutated in at least five independent Omicron sublineages that exhibited a high growth advantage [62,63].

The effect of nonadditive, epistatic relationships among RBD mutations was assessed using protein structure modeling by comparing the effects of all single mutants at the RBD-ACE2 interfaces for the Omicron variants, showing that structural constraints and stability requirements can drive virus evolution for a more complete antibody escape [64]. It was found that epistatic effects of mutations weakly destabilized antibody binding for Omicron variants relative to the ancestral background, thus suggesting that Omicron variants can display a higher potential for emergence of immune escape mutations than Delta or Wu-Hu-1 variants [64]. A systematic experimental analysis of the epistatic effects for the RBD residues using DMS approach in the Wu-Hu-1, Alpha, Beta, Delta, and Eta backgrounds showed that N501Y causes significant epistatic shifts in the mutational effects of Q498R and RBD residues 446-449 and 491-496 [65]. Q498R mutation can only marginally affect the ACE2 binding affinity [66] but in the background of N501Y mutant, the binding affinity can dramatically increase ([65,66] with Q498R and N501Y mutations together increasing the affinity by up to 387-fold compared to the Wu-Hu-1 strain [65]. As a result, it was suggested that the superior binding gain enabled by Q498R/N501Y double mutant may allow Omicron subvariants to accumulate immune escape mutations at other sites that are moderately destabilizing for ACE2 binding [67]. A systematic mapping of the epistatic interactions between the 15 BA.1 RBD mutations relative to the Wu-Hu-1 strain showed evidence of compensatory epistasis in which immune escape BA.1 mutations can individually reduce ACE2 binding but are compensated through epistatic couplings with affinity-enhancing mutations including Q498R and N501Y [67]. Recent evolutionary and functional studies revealed strong epistasis between pre-existing substitutions in BA.1/BA.2 variants and antibody resistance mutations acquired during selection experiments, suggesting that epistasis can lower the genetic barrier for antibody escape [68].

Computer simulations provided important atomistic and mechanistic advances into understanding the dynamics and function of the SARS-CoV-2 S proteins. All-atom molecular dynamics (MD) simulations of the full-length SARS-CoV-2 S glycoprotein embedded in the viral membrane, with a complete glycosylation profile provided detailed characterization of the conformational landscapes of the S proteins in the physiological environment [69-71]. Using distributed cloud-based computing, large scale MD simulations of the viral proteome observed dramatic opening of the S protein complex, predicting the existence of several cryptic epitopes in the S protein [72]. MD simulations of the S-protein in solution and targeted simulations of conformational changes between the open and closed forms revealed the key electrostatic interdomain interactions mediating the protein stability and kinetics of the functional spike states [73]. Using the replica-exchange MD simulations with solute tempering of selected surface charged residues, the conformational landscapes of the full-length S protein trimers were investigated, unveiling the transition pathways via inter-domain interactions, hidden functional intermediates along open-closed transition pathways and previously unknown cryptic pockets [74]. The enhanced MD simulations examined the stability of the S-D 614G variant showing that the mutation orders the 630-loop structure and allosterically alters global interactions between RBDs, forming an asymmetric and mobile down conformation and facilitating transitions toward up conformation [75]. Our previous studies revealed that the SARS-CoV-2 S protein can function as an allosteric regulatory machinery that is controlled by stable allosteric hotspots to modulate specific regulatory and binding functions [76-83]. By examining conformational landscapes and dynamic interaction networks in the SARS-CoV-2 Omicron structures, we found that Omicron mutational sites can be dynamically coupled through short and long-range interactions forming an adaptive allosteric network that controls balance and tradeoffs between conformational plasticity, protein stability, and functional adaptability [83].

A number of computational studies employed atomistic simulations and binding energy analysis to examine the interactions between the S-RBD Omicron and the ACE2 receptor. MD simulations of the Omicron RBD binding with ACE2 suggested that K417N, G446S, and Y505H mutations can decrease the ACE2 binding, while S447N, Q493R, G496S, Q498R, and N501Y mutations improve binding affinity with the host receptor [84]. By examining a large number of mutant complexes, it was found that high-affinity RBD mutations tend to cluster near known human ACE2 recognition sites supporting the view that combinatorial mutations in SARS-CoV-2 can develop in sites amenable for non-additive enhancements in binding and antibody evasion [85]. We examined differences in allosteric interactions and communications in the S-RBD complexes for Delta and Omicron variants using a combination of perturbation-based scanning of allosteric residue potentials and dynamics-based network analysis [86]. This study showed that G496S, Q498R, N501Y and Y505H correspond to the key binding energy hotspots and also contribute decisively to allosteric communications between S-RBD and ACE2. All-atom MD simulations of the RBD-ACE2 complexes for BA.1 BA.1.1, BA.2, an BA.3 Omicron subvariants were combined with a systematic mutational scanning of the RBD-ACE2 binding interfaces, revealing functional roles of the key Omicron mutational site R493, R498 and Y501 acting as binding energy hotspots, drivers of electrostatic interactions and mediators of epistatic effects and long-range communications [87].

In this study, we systematically examine the dynamics, stability and binding in the Omicron BA.2, BA.2.75, XBB.1 and XBB.1.5 RBD complexes with ACE2 using multiscale molecular simulations, in silico mutational scanning of the RBD residues for binding and stability and network-based community analysis of allosteric communications and epistatic interactions. The evolutionary and functional studies suggested that the emergence of the Omicron variants can be determined by multiple fitness trade-offs including the immune escape, binding affinity for ACE2, conformational plasticity, protein stability and allosteric modulation [78-80]. We employ MD simulations and network-based energetic analysis of RBD variants binding to quantify the balance and contributions of protein folding stability and binding interactions. By using network-based analysis of epistatic interactions, we examine a hypothesis that the emerging new variants, particularly XBB.1.5 may induce epistasis patterns where protein stability can promote evolvability by tolerating mutations in positions that confer beneficial phenotypes. Through the integration of computational approaches, we show that the enhanced RBD stability in the BA.2.75 and XBB.1.5 variants may be an important driving force for evolvability of new mutations and superior ACE2 binding. We argue that during the evolution of a virus, a mutation that helps the virus evade the human immune system might only be tolerated if the virus has acquired other mutations beforehand. This type of mutational interactions between stability hotspots and evolvability sites would constrain the evolution of the virus, since its capacity to take advantage of the second mutation depends on the first mutation having already occurred.

## Materials and Methods

### Structural modeling and refinement

The crystal structures of the BA.2 RBD-hACE2 (pdb id 7XB0), and BA.2.75 RBD-hACE2 complexes (pdb id 8ASY) were obtained from the Protein Data Bank [88]. During structure preparation stage, protein residues in the crystal structures were inspected for missing residues and protons. Hydrogen atoms and missing residues were initially added and assigned according to the WHATIF program web interface [89]. The missing loops in the studied cryo-EM structures of the SARS-CoV-2 S protein were reconstructed and optimized using template-based loop prediction approach ArchPRED [90]. The side chain rotamers were refined and optimized by SCWRL4 tool [91]. The protein structures were then optimized using atomic-level energy minimization with composite physics and knowledge-based force fields implemented in the 3Drefine method [92,93]. The refined structural models of the XBB.1 RBD-ACE2 and XBB.1.5 RBD-ACE2 complexes were obtained with the aid of MutaBind2 approach that utilizes molecular mechanics force fields and fast side-chain optimization algorithms via random forest (RF) method [94,95]. MutaBind2 utilizes FoldX approach [96,97] to introduce single or multiple point mutations on the crystal structure followed by robust side-chain optimization and multiple rounds of energy minimization using NAMD 2.9 program [98] with CHARMM36 force field [99].

### Coarse-Grained Brownian Dynamics Simulations

Coarse-grained Brownian dynamics (CG-BD) simulations have been conducted using the ProPHet (Probing Protein Heterogeneity) approach and program [100-103]. BD simulations are based on a high resolution CG protein representation where each amino acid is represented by one pseudo-atom at the Cα position, and two pseudo-atoms for large residues. The interactions between the pseudo-atoms are treated according to the standard elastic network model (ENM) in which the pseudo-atoms within the cut-off parameter, *R*_c_ = 9 Å are joined by Gaussian springs. The simulations use an implicit solvent representation via the diffusion and random displacement terms and hydrodynamic interactions through the diffusion tensor using the Ermak-McCammon equation of motions and hydrodynamic interactions as described in the original pioneering studies that introduced Brownian dynamics for simulations of proteins [104,105]. The stability of the SARS-CoV-2 S Omicron trimers was monitored in multiple simulations with different time steps and running times. We adopted Δ*t* = 5 fs as a time step for simulations and performed 100 independent BD simulations for each system using 100,000 BD steps at a temperature of 300 K. The CG-BD conformational ensembles were also subjected to all-atom reconstruction using PULCHRA method [106] and CG2AA tool [107] to produce atomistic models of simulation trajectories.

### All-Atom Molecular Dynamics Simulations

NAMD 2.13-multicore-CUDA package [98] with CHARMM36 force field [99] was employed to perform 500 ns all-atom MD simulations for each of the Omicron RBD-hACE2 complexes. The structures of the SARS-CoV-2 S-RBD complexes were prepared in Visual Molecular Dynamics (VMD 1.9.3) [108] by placing them in a TIP3P water box with 20 Å thickness from the protein. Assuming normal charge states of ionizable groups corresponding to pH = 7, sodium (Na+) and chloride (Cl-) counter-ions were added to achieve charge neutrality and a salt concentration of 0.15 M NaCl was maintained. All Na^+^ and Cl^-^ ions were placed at least 8 Å away from any protein atoms and from each other. The long-range non-bonded van der Waals interactions were computed using an atom-based cutoff of 12 Å with the switching function beginning at 10 Å and reaching zero at 14 Å. SHAKE method was used to constrain all bonds associated with hydrogen atoms. Simulations were run using a leap-frog integrator with a 2 fs integration time step. ShakeH algorithm of NAMD was applied for water molecule constraints. The long-range electrostatic interactions were calculated using the particle mesh Ewald method [109] with a cut-off of 1.0 nm and a fourth order (cubic) interpolation. Simulations were performed under NPT ensemble with Langevin thermostat and Nosé-Hoover Langevin piston at 310 K and 1 atm. The damping coefficient (gamma) of the Langevin thermostat was 1/ps. The Langevin piston Nosé-Hoover method in NAMD is a combination of the Nose-Hoover constant pressure method [110] with piston fluctuation control implemented using Langevin dynamics [111,112]. Energy minimization was conducted using the steepest descent method for 100,000 steps. All atoms of the complex were first restrained at their crystal structure positions with a force constant of 10 Kcal mol^-1^ Å^-2^. Equilibration was done in steps by gradually increasing the system temperature in steps of 20K starting from 10K until 310 K and at each step 1ns equilibration was done keeping a restraint of 10 Kcal mol-1 Å-2 on the protein C_α_ atoms. After the restraints on the protein atoms were removed, the system was equilibrated for an additional 10 ns. An NPT production simulation was run on equilibrated structures for 500 ns keeping the temperature at 310 K and constant pressure (1 atm).

### Distance Fluctuations Stability and Communication Analysis

We employed distance fluctuation analysis of the simulation trajectories to compute residue-based stability profiles. The fluctuations of the mean distance between each pseudo-atom belonging to a given amino acid and the pseudo-atoms belonging to the remaining protein residues were computed. The fluctuations of the mean distance between a given residue and all other residues in the ensemble were converted into distance fluctuation stability indexes that measure the energy cost of the residue deformation during simulations [100-103]. The distance fluctuation stability index for each residue is calculated by averaging the distances between the residues over the simulation trajectory using the following expression:

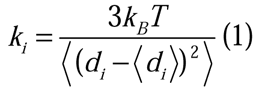

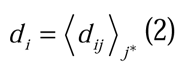

*d_ij_* is the instantaneous distance between residue *i* and residue *j*, *k_B_* is the Boltzmann constant, *T* =300K. 〈〉 denotes an average taken over the MD simulation trajectory and *d_i_* = 〈*d_ij_〉_j*_* is the average distance from residue *i* to all other atoms *j* in the protein (the sum over *j*_*_ implies the exclusion of the atoms that belong to the residue *i*). The interactions between the *C_α_* atom of residue *i* and the *C_α_* atom of the neighboring residues i-1 and *i*+1 are excluded in the calculation since the corresponding distances are constant. The inverse of these fluctuations yields an effective force constant *k_i_* that describes the ease of moving an atom with respect to the protein structure.

### Binding Free Energy Computations: Mutational Scanning and Sensitivity Analysis

The binding free energies were initially computed for the Omicron RBD-hACE2 complexes were performed for the crystal structures and the refined structural models using a contact-based predictor of binding affinity Prodigy [113-116]. We conducted mutational scanning analysis of the binding epitope residues for the SARS-CoV-2 S RBD-ACE2 complexes. Each binding epitope residue was systematically mutated using all substitutions and corresponding protein stability and binding free energy changes were computed. BeAtMuSiC approach [117-119] was employed that is based on statistical potentials describing the pairwise inter-residue distances, backbone torsion angles and solvent accessibilities, and considers the effect of the mutation on the strength of the interactions at the interface and on the overall stability of the complex. The binding free energy of protein-protein complex can be expressed as the difference in the folding free energy of the complex and folding free energies of the two protein binding partners:

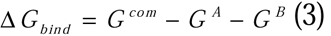

The change of the binding energy due to a mutation was calculated then as the following:

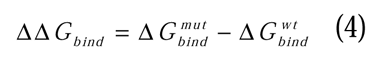

We leveraged rapid calculations based on statistical potentials to compute the ensemble-averaged binding free energy changes using equilibrium samples from simulation trajectories. The binding free energy changes were computed by averaging the results over 1,000 equilibrium samples for each of the studied systems.

### Network Community Analysis and Clique Model of Epistatic Interactions

A graph-based representation of protein structures [120,121] is used to represent residues as network nodes and the *inter-residue edges* to describe non-covalent residue interactions. The network edges that define residue connectivity are based on non-covalent interactions between residue side-chains. The residue interaction networks were constructed by incorporating the topology-based residue connectivity MD-generated maps of residues cross-correlations [122] and coevolutionary couplings between residues measured by the mutual information scores [123]. The edge lengths in the network are obtained using the generalized correlation coefficients *R_**MI**_ (x_**i**,_ x_**J**_*)associated with the dynamic correlation and mutual information shared by each pair of residues. The length (i.e. weight) *w_**ij**_ -- log[R_**MI**_ (x_**i**,_ x_**J**_)]* of the edge that connects nodes *i* and *j* is defined as the element of a matrix measuring the generalized correlation coefficient *R_**MI**_ (x_**i**,_ x_**J**_*)as between residue fluctuations in structural and coevolutionary dimensions. Network edges were weighted for residue pairs with *R_**MI**_ (x_**i**,_ x_**J**_)* > 0.5 in at least one independent simulation. The matrix of communication distances is obtained using generalized correlation between composite variables describing both dynamic positions of residues and coevolutionary mutual information between residues. Residue Interaction Network Generator (RING) program [124] was employed for generation of the residue interaction networks using the conformational ensemble where edges have an associated weight reflecting the frequency in which the interaction present in the conformational ensemble. The residue interaction network files in xml format were obtained for all structures using RING v3.0 webserver [124]. Network graph calculations were performed using the python package NetworkX [125]. Using the constructed protein structure networks, we computed the residue-based betweenness parameter. The short path betweenness of residue *i* is defined to be the sum of the fraction of shortest paths between all pairs of residues that pass through residue *i* :

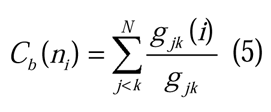

where *g_jk_* denotes the number of shortest geodesics paths connecting *j* and *k,* and *g _jk_* (*i*) is the number of shortest paths between residues *j* and *k* passing through the node *n_i_ .* Residues with high occurrence in the shortest paths connecting all residue pairs have a higher betweenness values. For each node *n,* the betweenness value is normalized by the number of node pairs excluding *n* given as (*N* -1)( *N* -2) / 2, where *N* is the total number of nodes in the connected component that node *n* belongs to. The normalized short path betweenness of residue *i* can be expressed as

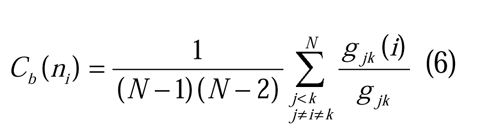

*g_jk_* is the number of shortest paths between residues *j* and k; *g _jk_* (*i*) is the fraction of these shortest paths that pass through residue *i*.

The Girvan-Newman algorithm [126] is used to identify local communities. In this approach, edge centrality (also termed as edge betweenness) is defined as the ratio of all the shortest paths passing through a particular edge to the total number of shortest paths in the network. The method employs an iterative elimination of edges with the highest number of the shortest paths that go through them. By eliminating edges, the network breaks down into smaller communities. The analysis of the interaction networks was done using network parameters such as cliques and communities. The *k* -cliques are complete sub graphs of size *k* in which each node is connected to every other node. In our application, a *k* -clique is defined as a set of *k* nodes that are represented by the protein residues in which each node is connected to all the other nodes. A *k* -clique community is determined by the Clique Percolation Method [127] as a subgraph containing *k* -cliques that can be reached from each other through a series of adjacent *k*-cliques. We have used a community definition according to which in a *k* -clique community two *k* - cliques share *k* −1 or *k* −2 nodes. Computation of the network parameters was performed using the Clique Percolation Method as implemented in the CFinder program [128]. Given the chosen interaction cutoff *^I^*_min_ we typically obtain communities formed as a union of *k* =3 and *k* =4 cliques. The interaction cliques and communities were considered to be dynamically stable if these interaction networks remained to be intact in more than 75% of the ensemble conformations. In the dynamic network model, it is assumed that allosteric interactions and long-range communications can be propagated through stable interaction networks in which the key network hubs serve as mediators of allosteric couplings. As dynamic couplings between RBD interface residues can be determined in simulations, we propose that strongly coupled residue positions may communicate and affect their ACE2 binding interactions via epistatic relationships. We assume that residues that belong to the same clique during simulations would have stronger dynamic and energetic couplings leading to synchronization and potentially epistatic effects. Using equilibrium ensembles and dynamic network modeling of the original RBD-ACE2 complexes for BA.2, BA.2.75, XBB.1 and XBB.1.5 variants, we use rapid mutational scanning to perturb modular network organization represented by a chain of inter-connected sable 3-cliques [129]. It is assumed that 3-clique community distributions may be perturbed during simulations and to examine the non-additive epistatic effect of a mutational site, we compared changes in the k-clique community distributions induced by different double mutations. Specifically, using the equilibrium ensembles we calculated the probability by which the two mutational sites belong to the same interfacial 3-clique. For this, we generated an ensemble of 1,000 protein conformation from 500 ns simulations of the studied RBD-ACE2 complexes. We computed the proportion *P_ab_* of snapshots in the ensemble in which the two mutational sites (*a*, *b*) belong to the interfacial 3-clique. *P_ab_* measures the probability that two sites (*a*, *b*) are kept in some 3-clique due to either direct or indirect interactions. The closer *P_ab_* is to 1, the more likely *a* and *b* tends to have a tight connection and potential local epistasis. By systematically using double mutational changes of the Omicron positions over the course of the MD simulation trajectory for the original RBD-ACE2 complexes we attribute mutational sites that belong to the same 3-clique to have local non-additive effects, while the effects of specific mutations on changes of the entire distribution and the total number of 3-cliques at the RBD-ACE2 interface will be attributed to long-range epistatic relationships.

## Results

### Structure-Based Analysis of Binding Interfaces and Intermolecular Contacts in the ACE2 Complexes with the Omicron RBD Variants

We first performed a comparative structural analysis of the Omicron RBD BA.2, BA.2.75, XBB.1 and XBB.1.5 complexes with the ACE2 receptor (Figure 1). Mutations G339D, S373P, S375F, K417N, N440K, S477N, T478K, E484A, Q493R, Q498R, N501Y, and Y505H in BA.2 are shared with the BA.1 variant, but BA.2 additionally carries S371F, T376A, D405N, and R408S mutations (Table 1, Figure 1A,B). BA.2.75 variant has nine additional mutations as compared to BA.2 in the NTD ( K147E, W152R, F157L, I210V, and G257S) and RBD (D339H, G446S, N460K, and R493Q) (Table 1, Figure 1C,D). G446S mutation is shared with Omicron BA.1, and R493Q reversed mutation is present in BA.4/BA.5 as well as in XBB.1 and XBB.1.5 subvariants. XBB.1 subvariant is a descendant of BA.2 and recombinant of BA.2.10.1 and BA.2.75 sublineages, featuring NTD mutations V83A, H146Q, Q183E, V213E, G252V and specific RBD mutations G339H, R346T, L368I, V445P, G446S, N460K, F486S, F490S and reversed R493Q (Table 1, Figure 1E,F). Importantly, some of these RBD mutations are known for their immune evasion functions, including R346T, G446S and F486S [130,131]. XBB.1.5 is essentially identical to XBB.1 with a critical single RBD modification F486P mutation (Figure 1E,F).

**Figure 1.**
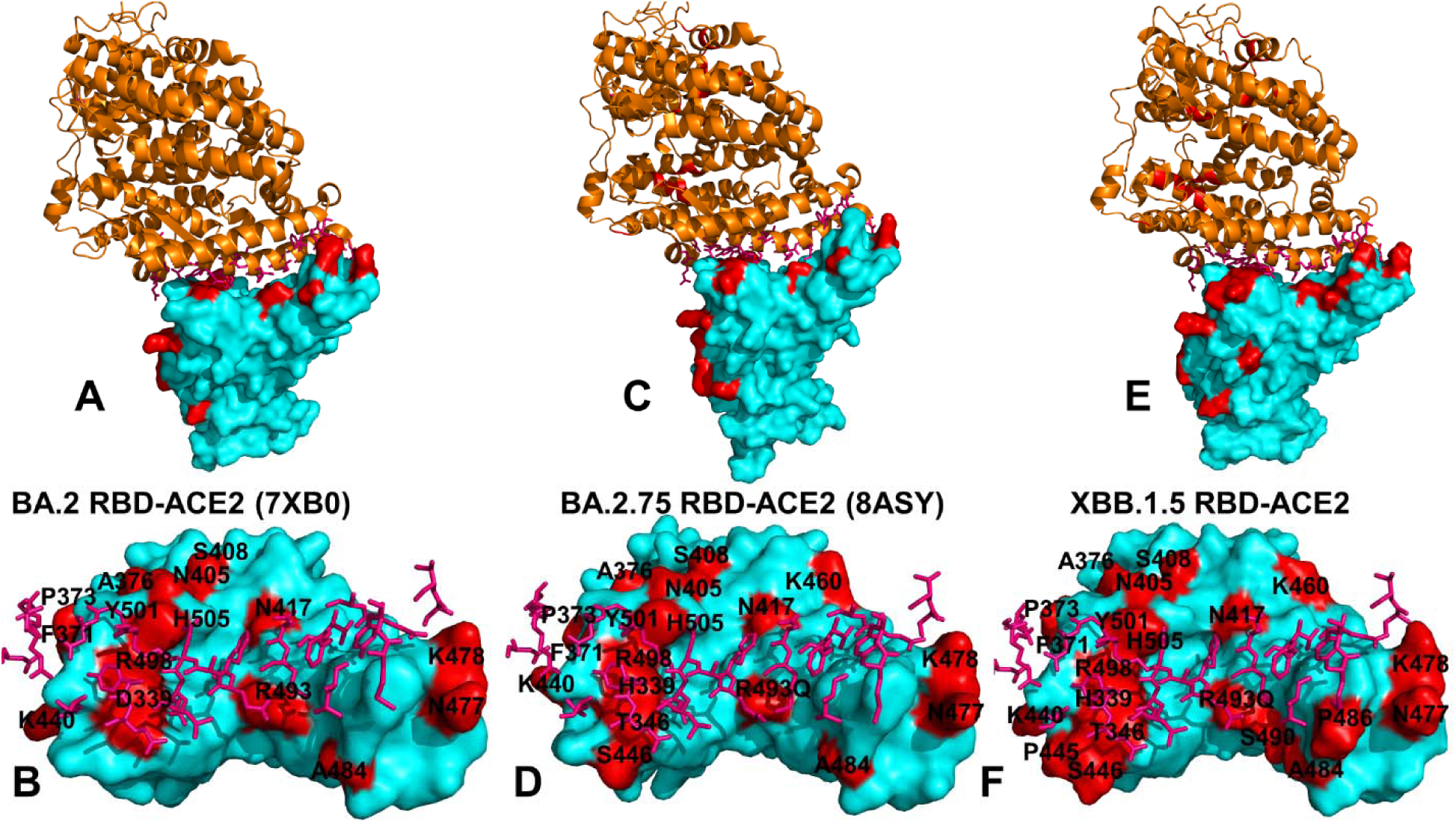
Structural organization of the SARS-CoV-2-RBD Omicron BA.2, BA.2.75, XBB.1 and XBB.1.5 complexes with human ACE2 enzyme. (A) The crystal structure of the Omicron RBD BA.2-ACE2 complex (pdb id 7XB0). The RBD is shown in cyan-colored surface and the bound ACE2 enzyme is in orange ribbons. (B) The RBD-BA.2 is shown in cyan surface from the top view. The ACE2 binding residues are shown in pink sticks. The Omicron RBD BA.2 sites (G339D, S371F, S373P, S375F, T376A, D405N, R408S, K417N, N440K, S477N, T478K, E484A, Q493R, Q498R, N501Y, Y505H) are shown in red surface and annotated. (C) The crystal structure of the Omicron RBD BA.2.75-ACE2 complex (pdb id 8ASY). The RBD is shown in cyan-colored surface and the bound ACE2 enzyme is in orange ribbons. (D) The RBD-BA.2.75 is shown from the top view. The ACE2 binding residues are shown in pink sticks. The Omicron RBD BA.2.75 sites (G339H, S371F, S373P, S375F, T376A, D405N, R408S, K417N, N440K, G446N, N460K, S477N, T478K, E484A, R493Q, Q498R, N501Y, Y505H) are shown in red surface and annotated. (E) The refined structural model of the Omicron RBD XBB.1.5-ACE2 complex. The RBD is shown in cyan-colored surface and the bound ACE2 enzyme is in orange ribbons. (F) The RBD-XBB.1 is shown from the top view and the ACE2 binding residues are shown in pink-colored sticks. The Omicron RBD XBB.1.5 sites (G339H, R346T, L368I, S371F, S373P, S375F, T376A, D405N, R408S, K417N, N440K, V445P, G446S,N460K, S477N, T478K, E484A, F486P, F490S, R493Q, Q498R, N501Y, Y505H) are shown in red-colored surface and annotated.

**Table 1.** Mutational landscape of the Omicron mutations for BA.1, BA.2, BA.275, XBB.1 and XBB.1.5 variants.

| Omicron Variant | Mutational landscape |
| --- | --- |
| BA.1 | A67, T95I, G339D, S371L, S373P, S375F, K417N, N440K, G446S, S477N, T478K, E484A, Q493R, G496S, Q498R, N501Y, Y505H, T547K, D614G, H655Y, N679K, P681H, N764K, D796Y, N856K, Q954H, N969K, L981F |
| BA.2 | T19I, G142D, V213G, G339D, S371F, S373P, S375F, T376A, D405N, R408S, K417N, N440K, S477N, T478K, E484A, Q493R, Q498R, N501Y, Y505H, D614G, H655Y, N679K, P681H, N764K, D796Y, Q954H, N969K |
| BA.2.75 | T19I, G142D, K147E, W152R, F157L, I210V, V213G, G257S, G339H, S371F, S373P, S375F, T376A, D405N, R408S, K417N, N440K, G446N, N460K, S477N, T478K, E484A, Q498R, N501Y, Y505H, D614G, H655Y, N679K, P681H, N764K, D796Y, Q954H, N969K |
| XBB.1 | T19I, V83A, G142D, Del144, H146Q, Q183E, V213E, G252V, G339H, R346T, L368I, S371F, S373P, S375F, T376A, D405N, R408S, K417N, N440K, V445P, G446S N460K, S477N, T478K, E484A, F486S, F490S, R493Q reversal, Q498R, N501Y, Y505H, D614G, H655Y, N679K, P681H, N764K, D796Y, Q954H, N969K |
| XBB.1.5 | T19I, V83A, G142D, Del144, H146Q, Q183E, V213E, G252V, G339H, R346T, L368I, S371F, S373P, S375F, T376A, D405N, R408S, K417N, N440K, V445P, G446S, N460K, S477N, T478K, E484A, F486P, F490S, R493Q |
|  | reversal, Q498R, N501Y, Y505H, D614G, H655Y, N679K,<br>P681H, N764K, D796Y, Q954H, N969K |

Mutations in F486 are of particular interest as F486V, F486I, F486S, F486P have been seen in other variants and arguably represent a convergent evolutionary hotspot shared by the recent wave of Omicron subvariants to optimize tradeoffs between binding affinity to ACE2 and immune evasion. According to the DMS experiments, among the most common F486 mutations (F486V/I/S/L/A/P), F486P imposes the lowest cost in RBD affinity loss and has the largest increase in RBD expression [58,59]. According to the escape calculator, F486 position is also one of the major hotspots for escaping neutralization by antibodies [132]. Structural analysis of the RBD complexes with ACE2 for BA.2 (Figure 1A,B) BA.2.75 (Figure 1C,D) and XBB.1/XBB.1.5 variants (Figure 1E,F) revealed very similar RBD conformations, the same binding mode of interactions with the host receptor and virtually identical topography of the binding interface.

A more scrupulous inspection of the RBDs with the mapped sites of the Omicron mutations highlighted subtle structural patterns of evolved mutations from BA.2 (Figure 1B) to BA.2.75 (Figure 1D) and XBB.1.5 variants (Figure 1F). First, the binding surface patch of Omicron mutations centered around the key binding hotpots R498,Y501 and H505 broadened in the BA.2.75 variant by adding T346 and S446 positions (Figure 1D) and expanded even further in the XBB.1.5 variant with T346, P445 and S446 sites (Figure 1F). One could also notice some consolidation of the RBD patch of mutational sites in the XBB.1.5 variant formed by the reversed R493Q, S490, A484, and P486 mutations that are linked to the flexible Omicron sites N477 and K478 (Figure 1F). For all variants, N440K is structurally disconnected from other Omicron sites, neither it is involved in direct intermolecular contacts with the ACE2. A similarly “isolated” and structurally peripheral N460K mutational site in BA.2.75 (Figure 1D) and XBB.1.5 (Figure 1F) is located away from the ACE2 binding interface.

Structural analysis of the binding interface residues and intermolecular contacts in the RBD-ACE2 complexes with the cutoff for the atom contact distance of 5.5 Å and cutoff for salt bridges at 3.5 Å [115,116] yielded an overall similar number of the RBD residues forming interaction contacts with ACE2 ( 30 RBD in the BA.2 RBD-ACE2 complex, 28 residues for BA.2.75 and 29 residues for XBB.1 and XBB.1.5 RBD complexes) (Table 2). The interaction atom pairs for the RBD-ACE2 complexes are listed in Tables S1-S4. Among instructive observations of this structural analysis was that mutational positions N440K and N460K are not involved in the intermolecular contacts with ACE2. It is evident that structure-based considerations alone are not sufficient to dissect the functional role of these Omicron mutations that in addition to their immune evading potential may have subtle effects on the RBD stability and ACE2 binding via long-range allosteric interactions. Structure-based binding free energy analysis using a contact-based Prodigy predictor of binding affinity [115,116] revealed a similar number of the interaction contacts mediated by charged residues as well as appreciable contributions of polar-nonpolar and nonpolar-nonpolar interactions (Table 2). The observed differences in the number of intermolecular contacts were fairly small and resulted in the predicted binding free energies that favored BA.2 and XBB.1.5 variants, showing a moderate loss in the binding affinity for XBB.1 (Table 2).

**Table 2.** The analysis of the interfacial residue-residue contacts and ensemble-averaged PRODIGY-based binding free energies for the Omicron RBD-hACE2 complexes.

| **Interactions** | **RBD**<br>**Omicron**<br>**BA.2-hACE2** | **RBD Omicron**<br>**BA.2.75-**<br>**hACE2** | **RBD**<br>**Omicron**<br>**XBB.1-**<br>**hACE2** | **RBD Omicron**<br>**XBB.1.5-**<br>**hACE2** |
| --- | --- | --- | --- | --- |
| Charged-charged | 8 | 4 | 5 | 5 |
| Charged-polar | 6 | 8 | 11 | 9 |
| Charged-apolar | 21 | 21 | 23 | 21 |
| Polar-polar | 6 | 5 | 7 | 7 |
| Polar-apolar | 20 | 18 | 22 | 18 |
| Apolar-apolar | 13 | 11 | 16 | 12 |
| $\Delta G$ computed(kcal/mol) | -12.1 | -11.4 | -11.3 | -12.4 |

Surprisingly, these simple computations correctly predicted a moderate decrease in the binding affinity of XBB.1 as compared to parental BA.2 [54]. Indeed, these experimental studies showed that the ACE2 binding affinities of XBB and XBB.1 exhibited a modest drop relative to that of BA.2 with K_d_ of 2.00 and 2.06 nM respectively compared to 0.95 nM for BA.2 [54]. A small loss in binding can be attributed to F486S mutation that could remove favorable hydrophobic interactions similar to what was observed for BA.4/5 variants where F486V and R493Q induced respectively a modest impairment and restoration of binding [58]. At the same time, structure-based binding affinity predictions failed to recognize the experimentally observed strongest binding of BA.2.75 immediately followed by XBB.1.5 variant [56]. Overall, this analysis suggested that structure-based assessments of the intermolecular interface contacts alone may be insufficient to fully capture subtle functional and binding affinity differences between the RBD-ACE2 complexes for different Omicron subvariants.

### Atomistic MD Simulations Reveal Common and Distinct Signatures of Conformational Dynamics and Interaction Patterns in the ACE2 Complexes with the Omicron RBD Variants

To examine the dynamic signatures of the Omicron variants, we conducted multiscale simulations of the RBD-ACE2 complexes that included multiple independent CG-BD simulations and atomistic reconstruction of the trajectories as well as all-atom MD simulations (Table 3, Figure 2). Through dynamics analysis, we probe the intrinsic conformational dynamics and identify differences in the RBD stability for BA.2, BA.2.75 and XBB subvariants. Using atomistic simulation analysis, we also examine a hypothesis whether some Omicron mutations may exert their effect on the stability and binding through long-range allosteric effects and epistatic relationships.

**Figure 2.**
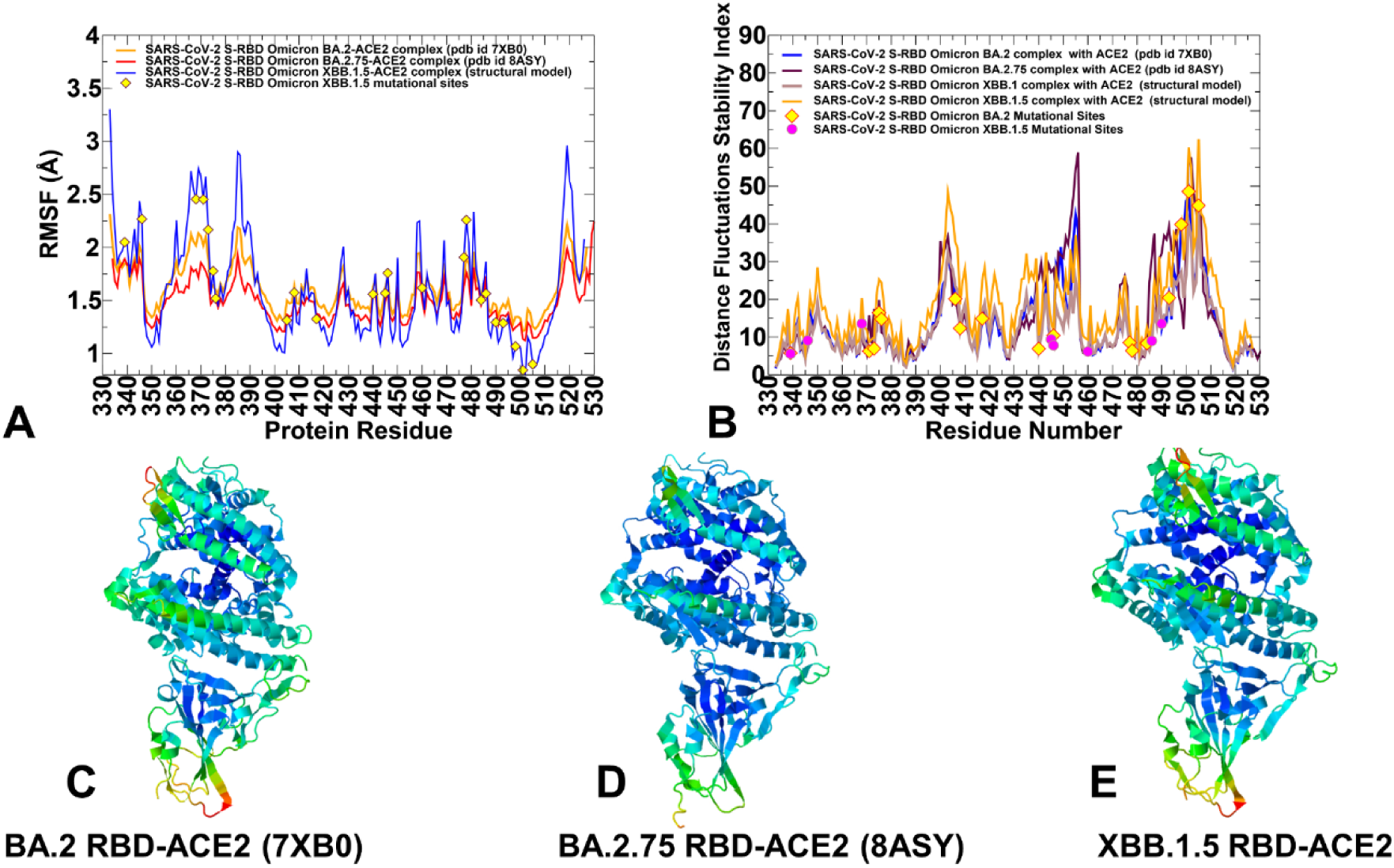
Conformational dynamics profiles obtained from all-atom MD simulations of the Omicron RBD BA.2, BA.2.75, XBB.1 and XBB.1.5 complexes with hACE2. (A) The RMSF profiles for the RBD residues obtained from MD simulations of the RBD BA.2-hACE2 complex, pdb id 7XB0 (in orange lines), RBD BA.2.75-hACE2 complex, pdb id 7XB0 (in red lines), and RBD XBB.1.5-hACE2 complex (in blue lines), The positions of Omicron RBD XBB.1.5 sites (G339H, R346T, L368I, S371F, S373P, S375F, T376A, D405N, R408S, K417N, N440K, V445P, G446S,N460K, S477N, T478K, E484A, F486P, F490S, R493Q, Q498R, N501Y, Y505H) are highlighted in yellow-colored filled circles. (B) The distance fluctuations stability index profiles of the RBD residues are shown for Omicron RBD BA.2 (in blue lines) BA.2.75 (in maroon lines), XBB.1 (in light brown lines) and XBB.1.5 (in orange lines). The positions of the Omicron BA.2 RBD sites (G339D, S371F, S373P, S375F, T376A, D405N, R408S, K417N, N440K, S477N, T478K, E484A, Q493R, Q498R, N501Y, Y505H) are highlighted in yellow-colored filled diamonds and XBB.1.5 mutational positions are depicted in magenta-colored circles. Structural maps of the conformational profiles are obtained from MD simulations of Omicron RBD variant complexes. Conformational mobility maps are shown for the Omicron RBD BA.2-hACE2 complex (C), the Omicron RBD BA.2.75-hACE2 complex (D), and the Omicron RBD XBB.1.5-hACE2 complex (E). The structures are shown in ribbons with the rigidity-flexibility sliding scale colored from blue (most rigid) to red (most flexible).

**Table 3.** Molecular simulations of the RBD-ACE2 complexes.

| **PDB** | **System** | **CG-BD** | **#simulations** | **All-atom**<br>**MD** |
| --- | --- | --- | --- | --- |
| 7XB0 | BA.2 RBD-hACE2 | 500,000<br>steps | 100 | 500 ns |
| 8ASY | BA.2.75 RBD-hACE2 | 500,000<br>steps | 100 | 500 ns |
| model | XBB.1 RBD-hACE2 | 500,000<br>steps | 100 | 500 ns |
| model | XBB.1.5 RBD-hACE2 | 500,000<br>steps | 100 | 500 ns |

Conformational dynamics profiles obtained from CG-BD and MD simulations were similar and revealed several important trends. Here, for clarity of presentations, we focused on all-atom MD simulations and analyzed the root mean square fluctuations (RMSF) distributions for the RBD residues (Figure 2A). The RBD has two subdomains, where the flexible RBM with a concave surface is involved in direct interaction contacts with hACE2 (Figure 1). The second subdomain is a five-stranded antiparallel β-sheet core region that functions as a stable core of the RBD. The conformational mobility distributions for the Omicron RBD complexe displayed several deep local minima corresponding to the RBD core residue cluster (residues 396-403) and the interfacial RBD positions involved in the contacts with the hACE2 receptor (residues 440-456 and 490-505 of the binding interface) (Figure 2A). The observed structural stability of the RBD core regions was also seen in our earlier simulation studies of the RBD Wu-Hu-1 and Omicron complexes [86,87], further confirming that these segments remain mostly rigid across all examined RBD complexes with hACE2. Noteworthy, the most stable RBD positions included several important hydrophobic stability centers F400, I402, F490, Y453, L455, A475, and Y489 (Figure 2A). Some of these hydrophobic RBD positions (Y453, L455, and Y489) are also involved in the favorable interfacial contacts with hACE2 and correspond to a stable conserved region of the RBD-hACE2 interface.

The RMSF profiles revealed signs of the greater stability for the BA.2.75 RBD as compared to other variants, featuring small thermal fluctuations not only in the ACE2-interacting sites but also only moderate displacements in the flexible RBD regions (residues 355-375 and 380-400) (Figure 2A). Despite similar dynamic profiles for all Omicron-hACE2 complexes we noticed that stable RBD core regions (residues 400-420, 430-450) exhibited even smaller fluctuations in the BA.2.75 and XBB.1.5 complexes (Figure 2A), suggesting the increased RBD stability for these variants which may be a relevant contributing factor of the stronger binding affinities seen for BA.2.75 and XBB.1.5 RBDs. The RMSF profile for XBB.1.5 is characterized by several deep minima corresponding to stabilized regions in the RBD core and particularly the key ACE2 binding interface cluster (residues 485-505) (Figure 2A). Furthermore, mapping of the Omicron XBB.1.5 mutational sites onto the dynamic profiles highlighted the increased stabilization of P486, S490, R493Q, R498, Y501 and H505 residues that become virtually immobilized in their interfacial positions (Figure 2A). Although this critical binding hotspot region is stable for all variants mainly due to strong interactions with ACE2, our findings indicated that the corresponding RBD positions become more rigid in BA.2.75 and XBB.1.5 (Figure 2A). This is generally consistent with the stronger binding affinities of these variants. The increased stabilization of the core RBD regions is accompanied by the moderately increased mobility localized around specific residues including RBM mutational sites N477 and K478. However, the majority of Omicron mutational sites in the studied RBD-ACE2 complexes maintained only a moderate degree of mobility with the exception of more flexible sites L368I, F371, N477 and K478 (Figure 2A).

It is worth noting that K440 and K460 mutational sites appeared to experience fairly moderate fluctuations, indicating that their intrinsic dynamic propensities may be partly curtailed in the BA.2.75 and XBB.1.5 RBD complexes to ensure greater folding stability of the RBD. The RBM residues that provide the contact interface with ACE2 also displayed relatively smaller movements and are largely stabilized in the complexes (Figure 2A). The mobile flexible RBM loops (residues 473-487) appeared to be partly constrained in the RBD-BA.2 and RBD-BA.2.75 complexes but remained more dynamic in XBB.1.5 (Figure 2A). Interestingly, a distal allosteric loop (residues 358–376) showed an appreciable mobility in the XBB.1.5 RBD as compared to BA.2 and BA.2.75 complexes, which may be potentially linked with the functional requirements for modulation of long-range couplings with the remote ACE2-binding interface residues (Figure 2A). In general, the RMSF profiles suggested that even though the distribution of rigid and flexible RBD regions is preserved and shared across all RBD complexes, the extent of rigidity and mobility in these regions may be modulated to elicit the increased rigidity of several stable RBD regions and maintaining local mobility of the flexible sites. Structural mapping of conformational dynamics profiles for BA.2 RBD (Figure 2B), BA.2.75 RBD (Figure 2C) and XBB.1.5 RBD (Figure 2D) complexes further illustrated a subtle but noticeable increase in the RBD rigidity for BA.2.75 and XBB.1.5 variants. Structural maps highlighted a progressive rigidification of the RBD-ACE2 binding interface regions in these complexes as well as the improved stability of the RBD core regions (Figure 2C,D). We argue that the improved RBD stability in BA.2.75 and XBB.1.5 complexes with ACE2 may be one of the factors linked with the experimentally observed enhancement in their binding affinity with ACE2.

Using conformational ensembles, we computed the fluctuations of the mean distance between each residue and all other protein residues that were converted into distance fluctuation stability indexes to measure the energetics of the residue deformations (Figure 2E). The high values of distance fluctuation indexes correspond to stable residues as they display small fluctuations in their distances to all other residues, while small values of this index would point to more flexible sites that experience larger deviations of their inter-residue distances. A comparative analysis of the residue-based distance fluctuation profiles revealed several dominant and common peaks reflecting similarity of the topological and dynamical features of the RBD-ACE2 complexes. The distributions showed that the local maxima for all RBD-ACE2 complexes are aligned with structurally stable and predominantly hydrophobic regions in the RBD core (residues 400-406, 418-421, 453-456) as well as key binding interface clusters (residues 495-505) that include binding hotspots R498 and Y501 (Figure 2E). Among RBD positions associated with the high distance fluctuations stability indexes are F400, I402, Y421, Y453, L455, F456, Y473, A475, and Y489 (Figure 2E). Despite a strikingly similar shape of the distributions for all Omicron RBD variants, which reflected the conserved partition of stable and flexible regions, the larger peaks were seen for BA.2.75 and XBB.1.5 RBD distributions (Figure 2E). This implies that the RBD core regions and ACE2-binding interface positions become progressively rigidified in the BA.2.75 and XBB.1.5 variants, suggesting the improved RBD folding stability and furter ehncement of the RBD-ACE2 binding interfaces for these variants. Importantly, common stability hotspots Y449, Y473, L455, F456, and Y489 are constrained by the requirements for the RBD folding and binding with the ACE2 host receptor, and therefore may be limited in evolving antibody escaping variants.

By highlghting BA.2 and XBB.1.5 mutational sites on the distriutions, it may be noticed that the majority of these RBD positions are characterized by low-to-moderate stability indexes, indicating a tendency of Omicron mutations to target conformationally adaptable regions in the RBD. Interestingly, XBB.1.5 mutational positions N440K, V445P, G446S, N460K, F486P and F490S displayed moderate distance fluctuation indexes, which may indicate presence of local mobility in these regions (Figure 2E). In contrary, the important RBD binding interface centers R498, Y501 and H505 featured high stability indexes, reflecting a considerable rigidification of these residues due to strong interactions with ACE2. According to our previous studies [78-83] residues with the high distance fluctuation indexes often serve not only as structurally stable centers but also as allosteric regulatory sites that control signal communication. Hence, Omicron sites R498, Y501 and H505 shared by all variants could function as key stability centers, binding hotspots as well as allosteric mediators of long-range communications in the RBD-ACE2 complexes (Figure 2). In this context, it may be instructive to relate our findings to the experimentally observed critical role of these hotspots in providing compensatory epistatic interactions with other Omicron sites [65-67] which allow to rescue sufficiently strong ACE2 binding and offset the effects of destabilizing immune escape mutations.

The observed “segregation” pattern of the Omicron sites portioning into a local cluster of structurally stable binding centers and a broadly distributed group of more flexible sites may reflect evolutionary requirements for energetically tolerant sites of immune evasion. The stability hotspots Y449, Y473, L455, F456, and Y489 featuring high indexes are constrained by the requirements for the RBD folding and binding with the ACE2 host receptor, and therefore may be limited in evolving antibody escaping mutations. At the same time, Omicron variant mutations in more dynamic sites may induce marginal destabilizing effects that are compensated for by localized cluster of structurally stable binding affinity hotspots. This interpretation is consistent with the notion that acquisition of functionally balanced substitutions to optimize multiple fitness tradeoffs between immune evasion, high ACE2 affinity and sufficient conformational adaptability may potentially be a common strategy of SARS-CoV-2 evolution executed by Omicron subvariants. Overall, the dynamic analysis of the RBD-ACE2 complexes suggested complementary roles of the Omicron mutation sites that may form a network of allosterically connected stable and more dynamic functional centers to enable modulation of structural stability, binding and long-range signaling.

### Computational Mutational Scanning of the RBD Residues Identifies Structural Stability and Binding Affinity Hotspots in the RBD-ACE2 Complexes : Quantifying Complementary Functional Effects of Omicron Mutations

Using the conformational ensembles obtained from MD simulations of the RBD-ACE2 complexes for BA.2, BA.2.75, XBB.1 and XBB.1.5 variants, we performed a systematic mutational scanning of the RBD residues in these complexes. In silico mutational scanning was done using BeAtMuSiC approach [117-119]. We enhanced this approach by averaging the binding free energy changes over the equilibrium ensembles. The reported binding free energy ΔΔG changes were evaluated by averaging the results of computations over 1,000 samples from MD simulation trajectories. The resulting “deep” mutational scanning heatmaps are reported for the RBD binding interface residues that make stable contacts with ACE2 in the course of simulations. To establish a relevance and validity of the in silico mutational scanning, we compared the results of the DMS experiments for the BA.1 and BA.2 RBD residues [58,59] with the computed mutational changes in the protein stability and binding for the BA.1 RBD-hACE2 (Figure 3A) and BA.2 RBD-hACE2 complexes (Figure 3B). A statistically significant correlation between the DMS experiments and mutational scanning data was observed, also highlighting the expected dispersion of the distributions. It is worth noting that the computed free energy changes reflected mutation-induced effects on both residue stability and binding interactions. Our findings are consistent with other simulation-based studies that showed correspondence between mutation-induced computed changes in the RBD stability and the experimental protein expression profiles [133]. It could be noticed that the computational predictions of destabilizing changes were often larger than the experimentally observed values. Nonetheless, the scatter plots showed a fairly appreciable correspondence between the predicted and experimental free energy differences, particularly for large destabilizing changes with ΔΔG > 2.0 kcal/mol (Figure 3A,B). This ensures an adequate identification of the major stabilization and binding affinity hotspots where mutations cause pronounced energetic changes. We also compared the DMS free energies for BA.2 RBD with the computationally predicted changes in XBB.1 and XBB.1.5 RBD complexes with ACE2 (Figure 3C,D).

**Figure 3.**
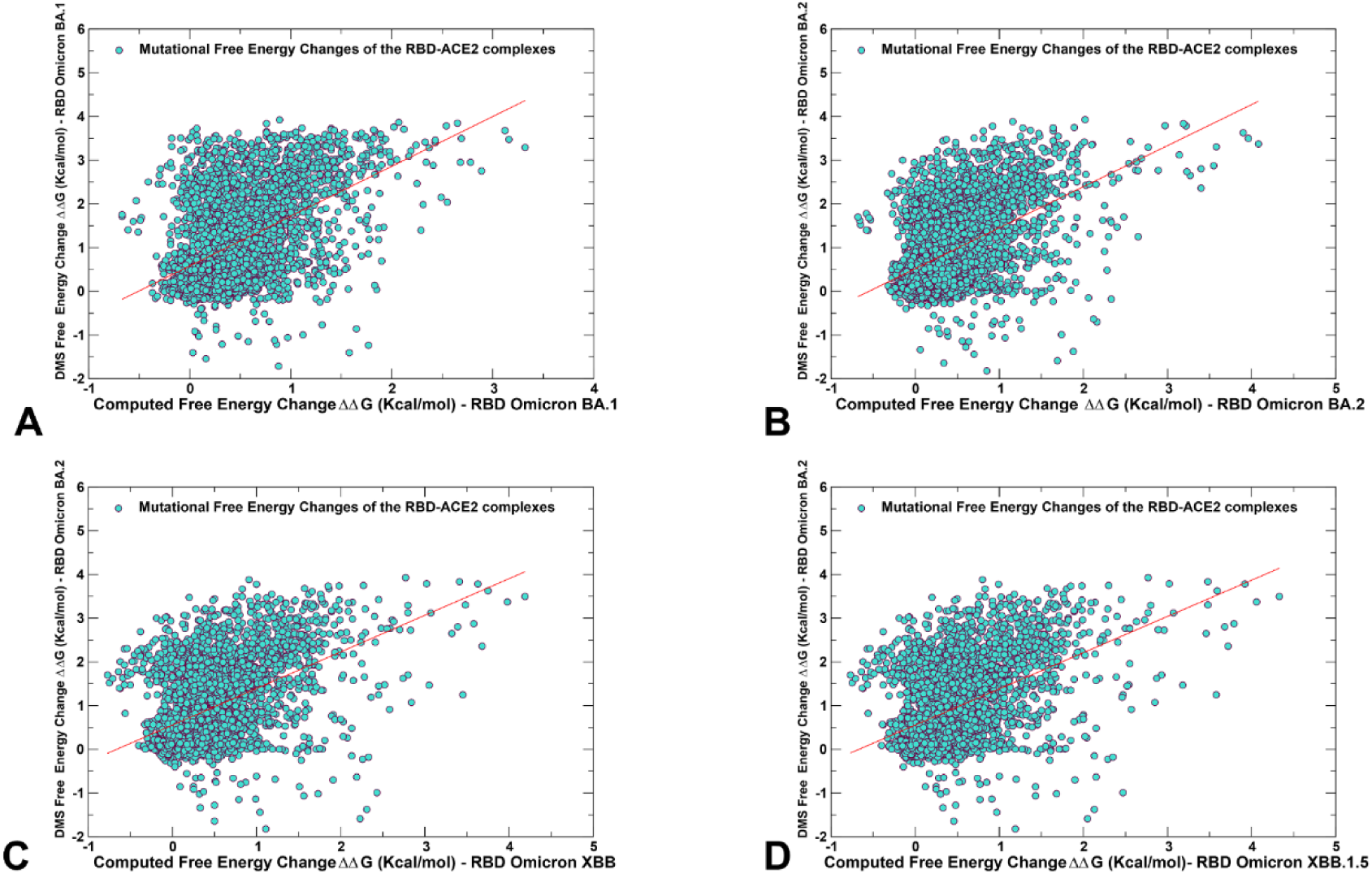
The scatter plots of the DMS-derived binding free energy changes for the RBD residues and computational mutational scanning of the RBD residues to estimate mutational effects on ACE2 binding. The effect on ACE2 receptor-binding affinity (Δ log10 *KD*) of every single amino-acid mutation in SARS-CoV-2 RBD was experimentally determined by high-throughput titration assays using DMS platform [58,59]. The results of computational mutational scanning of the RBD residues were averaged over conformational ensembles obtained from all-atom MD simulations. The scatter plot of the experimental and computed binding free energy changes from mutational scanning of the RBD residues in the Omicron RBD BA.1 complex, pdb id 7WBP (A) and RBD BA.2 complex, pdb id 7XB0 (B). The scatter plot of the experimental free energy changes from mutational scanning of the RBD residues in the RBD BA.2-ACE2 complex and computed binding free energy changes in XBB.1 RBD-ACE2 (C). The scatter plot of the experimental free energy changes from mutational scanning of the RBD residues in the RBD BA.2-ACE2 complex and computed binding free energy changes in XBB.1.5 RBD-ACE2 (D). The data points are shown in light-brown colored circles.

To analyze contributions of the RBD residues to protein stability, we also utilized a simplified SWOTein predictor which identifies the residue contributions to the global folding free energy through three types of database-derived statistical potentials that include inter-residue distances, backbone torsion angles and solvent accessibility [134,135]. According to this approach, positive folding free energy contributions indicate stability weaknesses while large negative folding free energy contributions for a given residue suggest stability strength for this position [134,135]. Despite its simplicity, this approach considers key contributions to the folding free energy associated with the enthalpic components, hydrophobic interactions and entropic estimations. The stability strengths and weaknesses are identified as residues that upon mutation result in strong destabilization or strong stabilization (Figure 4). The clusters of strong stability centers include several RBD regions including residues 400-405, 420-435, 453-458 and 477-489. Consistent with the dynamics analysis, among commonly shared folding stability centers are residues F400, I402, Y421, Y453, L455, F456, Y473, A475, and Y489 (Figure 4). Although the residue stability profiles are similar for all variants, we observed favorable folding stability values for a number of XBB.1.5 mutational sites including V445P, G446S,N460K, S477N, T478K, E484A, F486P, F490S, R493Q, Q498R, N501Y, Y505H (Figure 4D). Of particular significance is a strong stability peak associated with F486P position in XBB.1.5 variant (Figure 4D), thus indicating that convergent mutations in this position can modulate RBD stability and affect ACE2 binding. These results suggested that XBB.1.5 mutations may improve the RBD folding stability and concurrently enhance the ACE2 binding affinity compared to other examined variants through both the improved local contacts and globally by promoting a more stable RBD binding interface.

**Figure 4.**
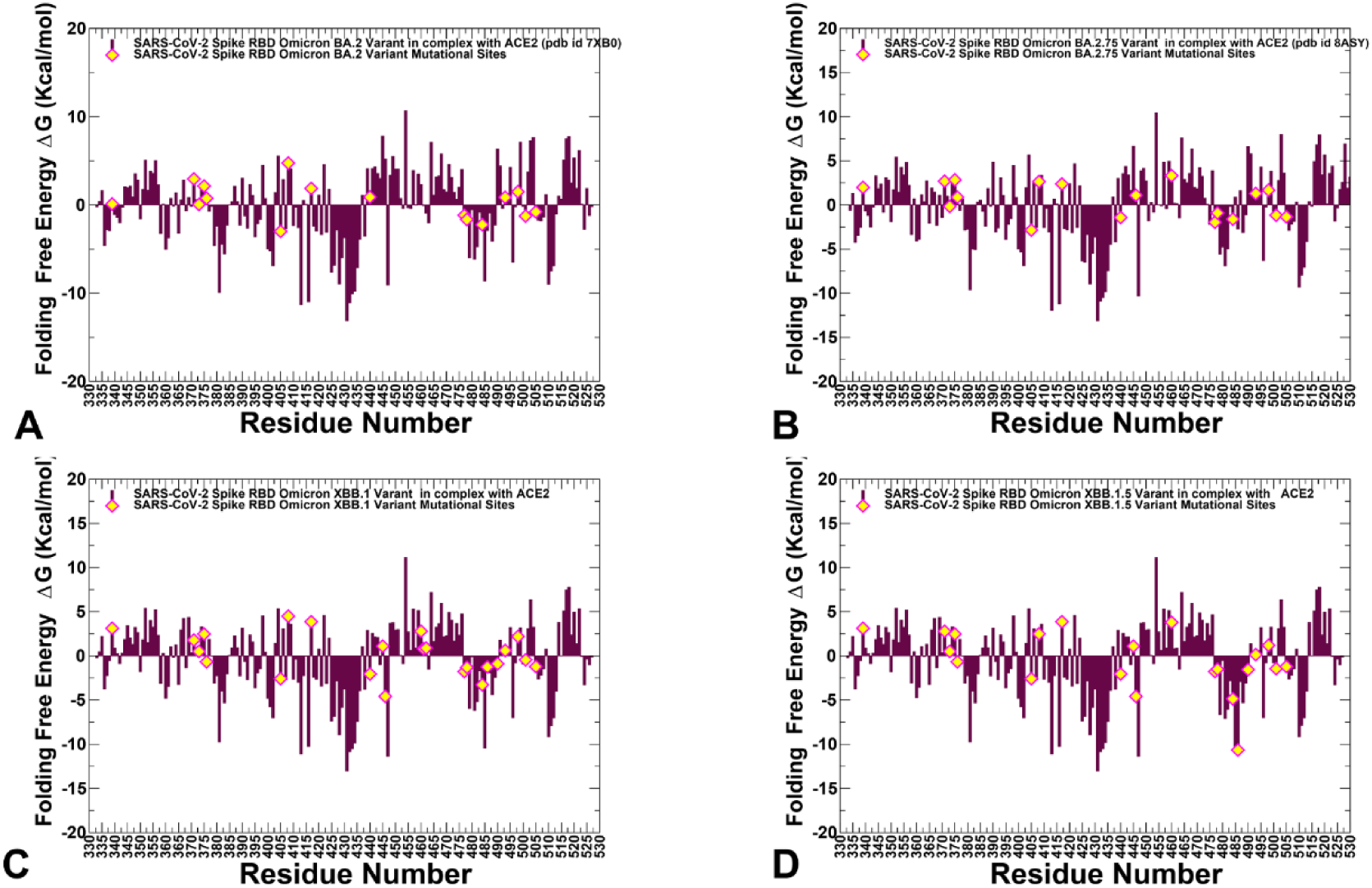
The folding free energies of the RBD residues for the SARS-CoV-2 S Omicron RBD-hACE2 complexes for BA.2, BA.2.75, XBB.1 and XBB.1.5 variants. The folding free energies for the RBD residues in the BA.2 RBD-hACE2 complex (A), BA.2.75 RBD-hACE2 complex (B), XBB.1 RBD-hACE2 complex (C) and XBB.1.5 RBD-hACE2 complex (D). The profiles are shown in maroon-colored bars. The positions of the Omicron mutational sites on the profiles are highlighted in yellow-colored filled diamonds. Note that the XBB.1.5 mutational sites V445P, G446S,N460K, S477N, T478K, E484A, F486P, F490S, R493Q, Q498R, N501Y, Y505H showed favorable stability, particularly evident for F486P mutation.

To provide a systematic comparison, we constructed mutational heatmaps for the RBD binding interface residues (Figure 5). Consistent with the DMS experiments [58,59], the strongest binding energy hotspots in BA.2 and BA.2.75 complexes corresponded to hydrophobic residues Y453, F456, Y473, Y489 and Y501 that play a decisive role in binding for all Omicron complexes (Figure 5A,B). Mutational heatmaps illustrated that the majority substitutions in these key interfacial positions can lead to a considerable loss in the RBD folding stability and binding affinity with ACE2. Interestingly, these positions are distributed across both major patches of the binding interface. This analysis is also consistent with our previous studies, suggesting that these conserved hydrophobic RBD residues may be universally important for binding across all Omicron variants and act as stabilizing sites of the RBD stability and binding affinity [86,87]. The common energetic hotspots Y453, F456, Y489 and Y501 found in the computational mutational scanning also emerged as critical stability and binding hotspots in the DMS studies [58,59]. In addition, mutational scanning of the RBD residues F486, N487 and H505 showed appreciable and consistent destabilization changes, placing these residues as second group of the energetic centers (Figure 5).

**Figure 5.**
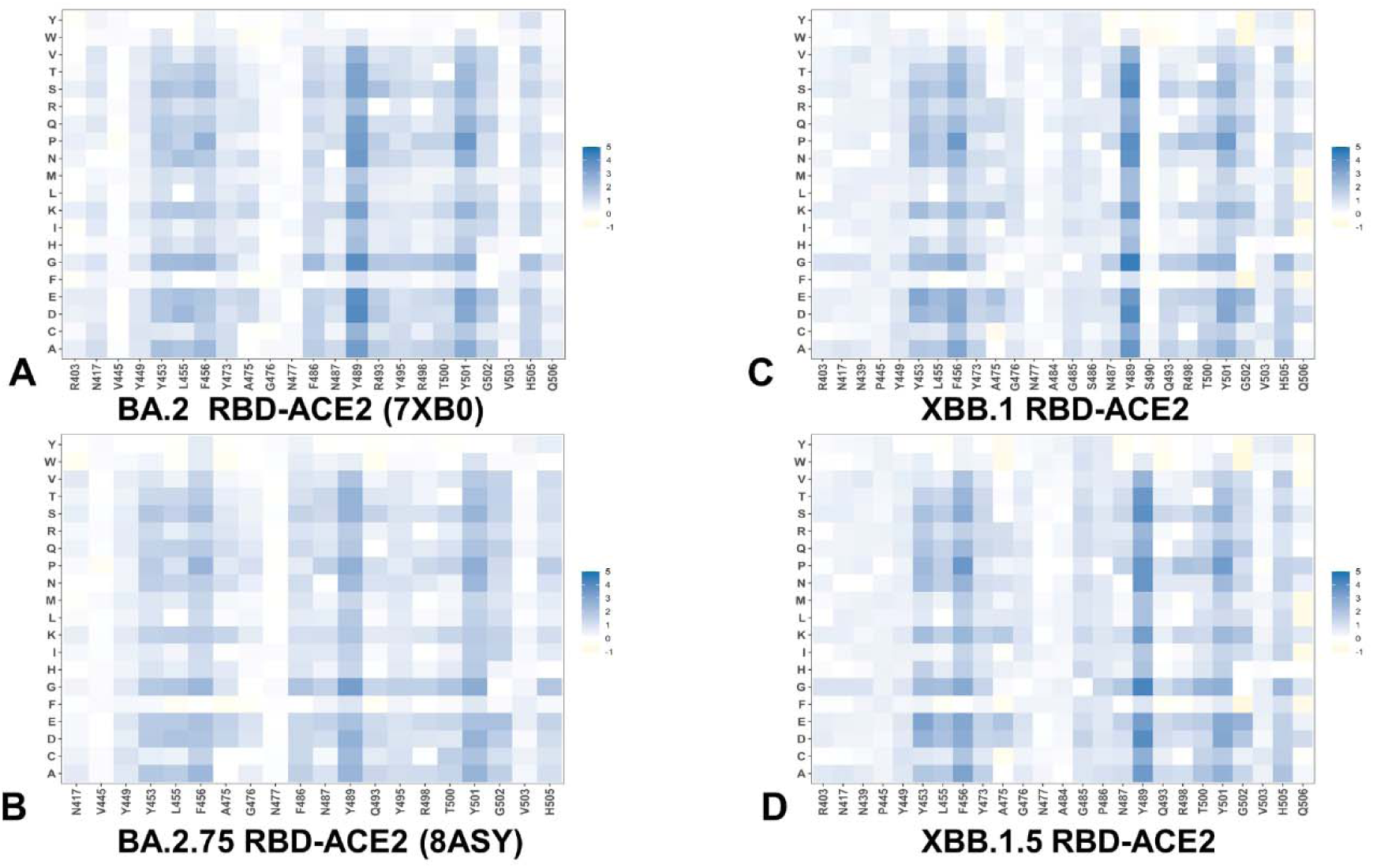
Ensemble-based dynamic mutational profiling of the RBD intermolecular interfaces in the Omicron RBD-hACE2 complexes. The mutational scanning heatmaps are shown for the interfacial RBD residues in the Omicron RBD BA.2-hACE2 (A), Omicron RBD BA.2.75-hACE2 (B), Omicron RBD XBB.1-hACE2 (C), and Omicron RBD XBB.1.5-hACE2 complexes (D). The heatmaps show the computed binding free energy changes for 20 single mutations of the interfacial positions. The standard errors of the mean for binding free energy changes were based on MD trajectories and selected samples (a total of 1,000 samples) are within ∼ 0.18-0.22 kcal/mol using averages from MD trajectories.

The computed heatmaps are consistent with the DMS experiments in which G446S and F486V mutations decrease ACE2 affinity of BA.2 (by -0.1 & -0.5 log10 Kd, respectively), while R493Q buffers these mutations by slightly increasing ACE2 affinity [58,59]. In the context of a comparative analysis between BA.2, BA.2.75, XBB.1 and XBB.1.5 variants, it is particularly relevant to notice that F486 position which is highly favorable for the RBD stability and binding is mutated to S486 in XBB.1 (Figure 5C) and P486 in XBB.1.5 (Figure 5D). The predicted binding free energy changes showed that P486 position of XBB.1.5 (Figure 5D) is less tolerant to modifications and more energetically favorable for the RBD stability and binding as compared to S486 in XBB.1 (Figure 5C). Importantly, mutational scanning revealed an appreciably destabilizing free energy change ΔΔG = 0.78 kal/mol for P486S modification in the XBB.1.5 structure, while the reversed S486P in XBB.1 yielded a modest favorable change with ΔΔG = -0.2 kal/mol (Figure 5C,D). These results agree with the experiments, indicating that F486P mutation in XBB.1.5 may rescue the loss of binding affinity in XBB.1. The emergence of F486V/S/ mutations is mainly due to their beneficial role for antibody escape at the expense of ACE2 affinity. Our results provided supporting evidence to the notion that the key functional difference between XBB.1.5 and its immediate parent XBB.1 is that XBB.1.5 has traded the costly F486S mutation for a more favorable F486P mutation thus enhancing both RBD stability and ACE2 binding. We argue that a synergistic effect of the restored binding and the improved RBD stability may favor transmissibility and the observed surge of the XBB.1.5 variant. Indeed, according to functional experiments F486P imposes the lowest cost in RBD affinity loss and has the largest increase in RBD expression [58,59].

Mutational scanning heatmaps also highlighted an interesting trend which may be relevant for the evolutionary role of the R493Q reversion which is in BA.2.75/XBB.1/XBB.1.5 as well as in BA.4/5. This mutation is reversion of Q493R that occurred early in BA.2’s evolution and while R493Q is not a major antigenic mutation it can arguably enable both F486V (in BA.4/5) and G446S (in BA.2.75) [58,59]. According to mutational scanning results, R493Q mutation is favorable for ACE2 binding with ΔΔG = -0.41 kal/mol in XBB.1 and with ΔΔG = -0.24 kal/mol in XBB.1.5 (Figure 5C,D). This is consistent with the experimental evidence which emphasized the fact that F486S/P mutations may have been selected to promote immune escape and are buffered by the favorable binding induced by the reversed R493Q change [58,59]. A similar effect was observed for F490S mutation that is marginally unfavorable for binding showing that the reversed S490F modification would improve binding with ΔΔG = -0.37 kcal/mol (Figure 5). Our data provide strong support to the emerging mechanism suggesting that XBB lineage may have evolved to evade immune suppression and outcompete other Omicron subvariants through mutations of F486 which is a hotspot for establishing protective immunity against the virus. To provide a better tradeoff between virus fitness requirements, XBB.1.5 acquired F486P which partly restored loss in ACE2 binding acting cooperatively with R493Q, R498 and Y501 while retaining the immune escape potential [136].

We also report mutational heatmaps focused only on the Omicron variant mutations in BA.2, BA.2.75, XBB.1 and XBB.1.5 RBD complexes (Figure 6) providing the estimate of the aggregate mutational impact of these variations on both RBD stability and ACE2 binding. The maps for BA.2 and BA.2.75 displayed a clear segregation between commonly shared binding hotspot sites (R493/Q493, R498, Y501, H505) and remaining mutational positions showing a notable tolerance to modifications (Figure 6A,B) thus allowing for evolution in some of these sites without sacrificing the RBD stability or ACE2 binding. A similar pattern of the binding free energy changes was seen in mutational scanning maps for XBB.1 and XBB.1.5 RBD residues (Figure 6C,D). Notably, for XBB.1.5 RBD, we observed an additional small group of Omicron sites (I368, N417, P445 and P486) that are more sensitive to modifications, but other Omicron positions are tolerant to substitutions producing a small effect on the RBD stability and binding (Figure 6D). Together with the mutational heatmaps for the RBD binding interface residues, these findings showed the presence of several folding stability centers (Y453, F456, Y489) and binding affinity hotspots (Q493, R498, T500, N501Y), while the remaining sites can potentially tolerate functionally beneficial destabilizing mutations.

**Figure 6.**
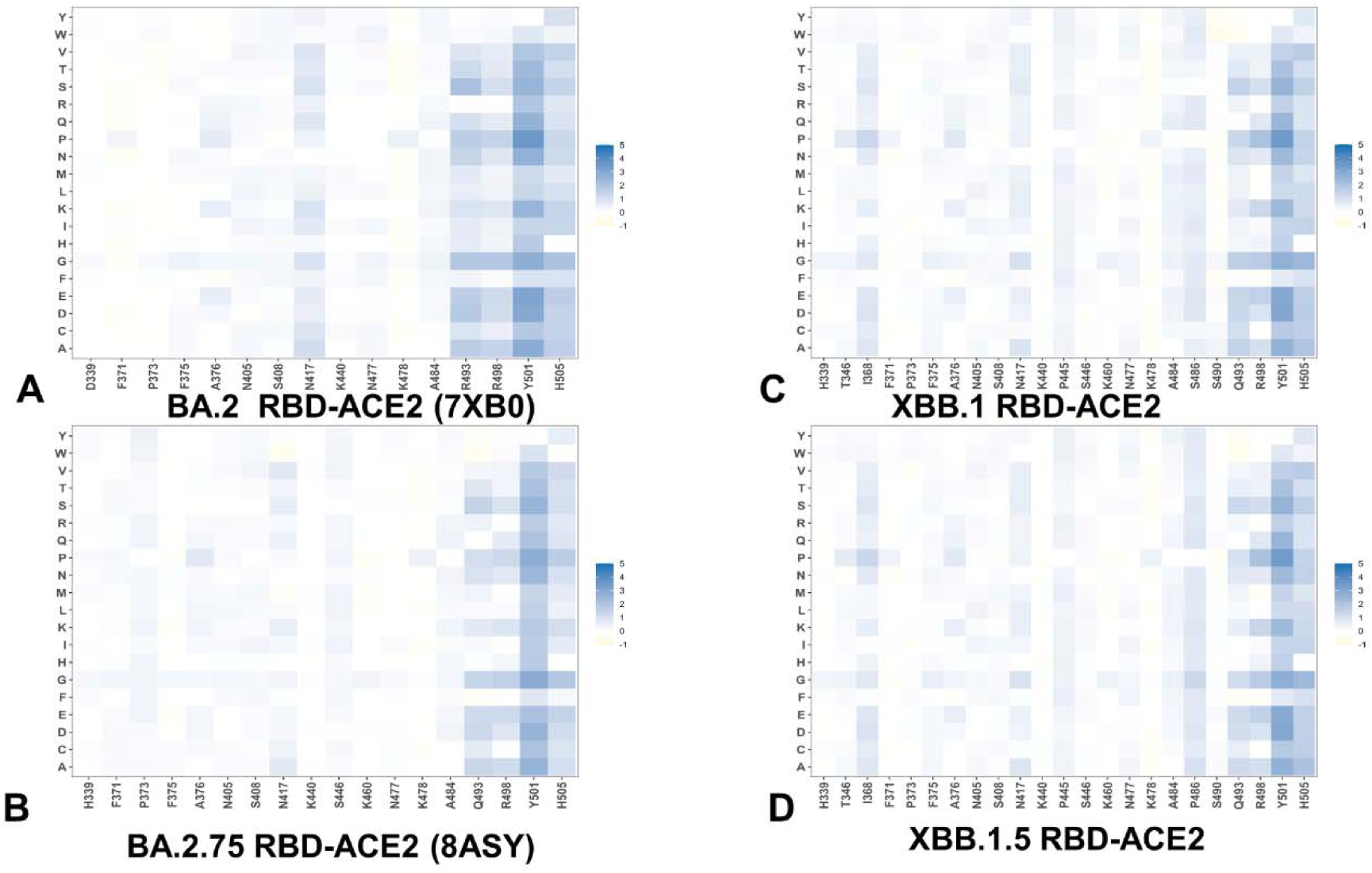
Ensemble-based dynamic mutational profiling of the Omicron RBD mutational sites in the RBD-hACE2 complexes. The mutational scanning heatmaps are shown for the BA.2 mutational sites (G339D, S371F, S373P, S375F, T376A, D405N, R408S, K417N, N440K, S477N, T478K, E484A, Q493R, Q498R, N501Y, Y505H) in the Omicron RBD BA.2-hACE2 complex (A). Mutational scanning heatmap for the BA.2.75 mutational sites (G339H, S371F, S373P, S375F, T376A, D405N, R408S, K417N, N440K, G446N, N460K, S477N, T478K, E484A, R493Q, Q498R, N501Y, Y505H) in the Omicron RBD BA.2.75-hACE2 complex (B). Mutational heatmap for the XBB.1 mutational sites (G339H, R346T, L368I, S371F, S373P, S375F, T376A, D405N, R408S, K417N, N440K, V445P, G446S,N460K, S477N, T478K, E484A, F486S, F490S, R493Q, Q498R, N501Y, Y505H) in the Omicron RBD XBB.1-hACE2 complex (C). Mutational heatmap for the XBB.1.5 mutational sites (G339H, R346T, L368I, S371F, S373P, S375F, T376A, D405N, R408S, K417N, N440K, V445P, G446S,N460K, S477N, T478K, E484A, F486P, F490S, R493Q, Q498R, N501Y, Y505H) in the Omicron RBD XBB.1.5-hAC2 (D). The standard errors of the mean for binding free energy changes were based on MD trajectories and selected samples (a total of 1,000 samples) are within ∼ 0.18-0.22 kcal/mol using averages from MD trajectories.

Structural mapping of the RBD binding interface epitopes, hot spot residues and the Omicron mutational sites (Figure 7) highlighted considerable similarities in the topography of the binding epitopes and “expansionary” character of the XBB.1 and XBB.1.5 variant positions (Figure 7E-H). The central binding interface cluster anchored by R498 and Y501 hotspots becomes further consolidated with the addition of T346, P445 and S446 mutations (Figure 7G,H). This key binding interface region includes sies of Omicron mutations R498, Y501 and H505 that provide the bulk of the ACE2 binding affinity. A visual inspection of these structural maps pointed to a denser Omicron mutational “shield” for XBB.1 (Figure 7E,F) and XBB.1.5 (Figure 7G,H) that encircles the core of the RBD epitope and partially overlaps with its peripheral areas. The maps further illustrated the notion that majority of the Omicron mutations, with the exception of R498/Y501/H505 cluster, tend to emerge near borders of the binding epitope causing relatively moderate changes in RBD stability and binding while targeting positions to induce a broad antibody escape (Figure 7).

**Figure 7.**
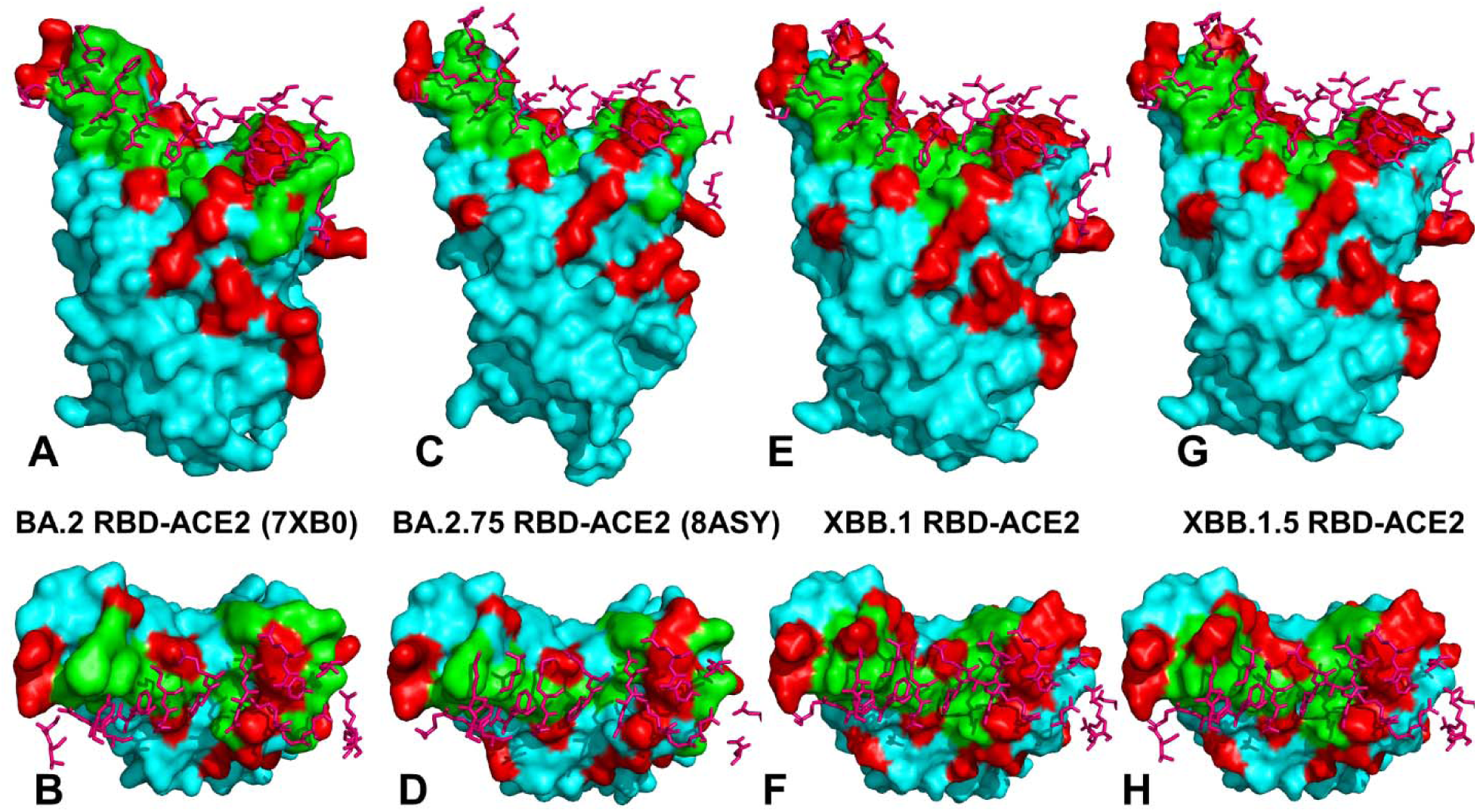
Structural mapping of the RBD binding epitopes of the SARS-CoV-2-RBD Omicron BA.2, BA.2.75, XBB.1 and XBB.1.5 complexes with human ACE2 enzyme. (A) The crystal structure of the Omicron RBD BA.2-ACE2 complex (pdb id 7XB0). The RBD binding epitope is shown in green-colored surface. The ACE2 binding residues are in pink sticks. The Omicron RBD BA.2 sites are shown in red surface. (B) The top view of the BA.2 RBD-ACE2 complex with the binding epitope residues in green and BA.2 mutations in red. (C,D) The crystal structure of the Omicron RBD BA.2.75-ACE2 complex (pdb id 8ASY). The RBD-BA.2.75 binding epitope (in green), the ACE2 binding residues (pink sticks) and BA.2.75 RBD sites (in red) are shown. (E,F) The modeled structure of the Omicron RBD XBB.1-ACE2 complex. The RBD-XBB.1 binding epitope (in green), the ACE2 binding residues (pink sticks) and XBB.1 RBD sites (in red) are shown. (G,H) The modeled structure of the Omicron RBD XBB.1.5-ACE2 complex. The RBD-XBB.1.5 binding epitope (in green), the ACE2 binding residues (pink sticks) and XBB.1.5 RBD sites (in red) are shown.

We report a more detailed residue-based binding free energy profile, focusing on the Omicron mutational changes in the RBD-ACE2 complexes for BA.2, BA.2.75, XBB.1 and XBB.1.5 variants (Figure 8). Overall, the profiles revealed a similar trend across all studied variants. In the BA.2 variant, Omicron mutations G339D, S371F, S373P, S375F, T376A, D405N, R408S, N440K, S477N, T478K, E484A are essentially stability-neutral, having only a marginal effect on binding (Figure 8A). K417N may induce a moderate change in ACE2 binding while promoting the increased neutralization escape potential of the Omicron variant from antibodies. At the same time Q498R, N501Y and Y505H seem to incur more significant stabilizing changes. Notably, and consistent with the experiments, N501Y modification showed a markedly larger stabilizing change in the ACE2 binding. The dominant stabilizing effect of N501Y is even more significant for BA.2.75 variant with R493Q, Q498R and Y505H also contributing appreciably to the enhanced binding affinity (Figure 8B).

**Figure 8.**
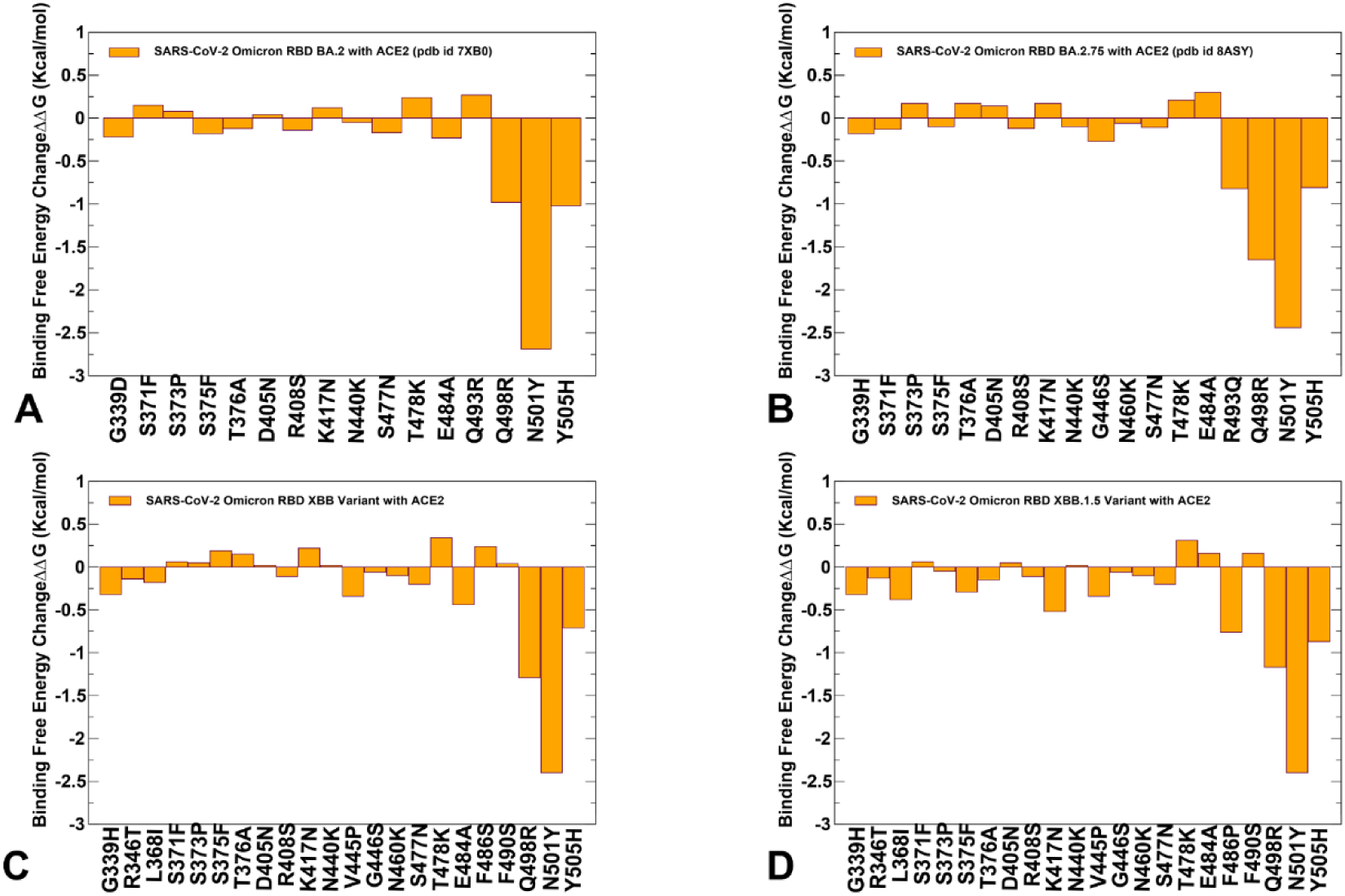
The predicted binding free energy changes in the RBD BA.2-hACE2 complex for BA.2 RBD mutations G339D, S371F, S373P, S375F, T376A, D405N, R408S, K417N, N440K, S477N, T478K, E484A, Q493R, Q498R, N501Y, Y505H (A). The computed binding free energy changes in the RBD BA.2.75-hACE2 complex for BA.2.75 RBD mutations G339H, S371F, S373P, S375F, T376A, D405N, R408S, K417N, N440K, G446N, N460K, S477N, T478K, E484A, R493Q, Q498R, N501Y, Y505H (B). The computed binding free energy changes in the RBD XBB.1-hACE2 complex for XBB.1 RBD mutational sites G339H, R346T, L368I, S371F, S373P, S375F, T376A, D405N, R408S, K417N, N440K, V445P, G446S,N460K, S477N, T478K, E484A, F486S, F490S, R493Q, Q498R, N501Y, Y505H (C). The binding free energy changes in the RBD XBB.1.5-hACE2 complex for XBB.1.5 RBD mutations G339H, R346T, L368I, S371F, S373P, S375F, T376A, D405N, R408S, K417N, N440K, V445P, G446S,N460K, S477N, T478K, E484A, F486P, F490S, R493Q, Q498R, N501Y, Y505H (D).

While the trend of most mutations being largely neutral for binding persists for all examined Omicron variants, we observed that for RBD XBB.1.5 the appreciably favorable binding free energy changes are associated with F486P, R493Q reversed mutation, Q498R, N501Y and Y505H mutations (Figure 8D). In addition, the other XBB.1.5 mutations are largely neutral, highlighting the tolerance of the respective positions to modifications. Consistent with the experimental data [65,67], these profiles further exemplified that functionally beneficial effects of destabilizing mutations (such as immune escaping potential and conformational adaptability) may be contingent on compensatory effects provided by a binding hotspot cluster centered on the R498/Y501 positions. These findings highlighted the important feature of a mechanism that may be characterized by the stability hotspots that ensure sufficient RBD stability and a spatially localized group of key binding affinity centers, while allowing for functionally beneficial but binding neutral (or moderately destabilizing) mutations in other positions to balance tradeoffs between immune evasion and ACE2 binding. The revealed patterns are reminiscent of direct evolution studies showing that enhanced protein stability in key sites can promote broader evolvability and expand a range of beneficial mutations while retaining the stability of the protein fold [137,138]. The related studies further elaborated that epistatic interactions between protein sites are mediated by stability, and that stabilizing mutations are often pre-requisites for adaptive destabilizing substitutions [139]. Moreover, these fascinating evolutionary studies suggested that stabilizing mutations promote protein evolvability and tolerance to destabilizing mutations that often contribute to immune escape. The presented mutational heatmaps for Omicron mutational sites are generally consistent with this mechanism, revealing a small group of shared stabilizing RBD positions that protect stability and binding affinity allowing for substantial evolvability at the remaining tolerant sites.

We argue that by expanding the range of stability/binding affinity hotspots and recruiting F486P/R493Q positions to maintain stability and restore binding affinity, XBB.1.5 variant can optimize both ACE2 binding and immune escaping profiles to achieve a superior viral fitness. These findings may be relevant in the context of epistasis in which nonlinear couplings between mutations may determine balance and tradeoffs between stability, evolvability and functions. We suggest that a balance between protein stability requirements and ACE2 binding affinity can promote evolvability of XBB variants by tolerating mutations in positions that could confer beneficial phenotypes. Based on this assertion, we also propose that Omicron mutational effects are mediated by protein stability and that the individually destabilizing RBD mutations may be counterbalanced via allostery and epistasis by stabilizing and affinity-enhanced mutations.

### Dynamic-Based Network Modeling and Community Analysis of the Omicron RBD-ACE2 Complexes Detail Allosteric Role of the Binding Energy Hotspots and Epistatic Relationships of the RBD Residues

Functional studies [65-67] suggested that evolutionary potential of the Omicron variants could be enhanced through epistatic interactions between variant mutations and the broader mutational repertoire available to the S proteins. In particular, it was suggested that weak epistasis in the Wu-Hu-1 original strain may become much stronger in newly emerged the Omicron variants as a potential virus mechanism to counteract structural limitations and stability constraints for continued evolution [67]. Here, we employed the ensemble-based modeling of the residue interaction networks utilizing a graph-based description [120,121] in which both dynamics [122] and coevolutionary couplings between protein residues [123] determine the strength of the interaction links. Using the dynamic inter-residue interaction networks, we performed community decomposition and characterized stable local interaction modules in which residues are densely interconnected through coupled interactions and dynamic correlations (Figure 9). A community-based model of allosteric interactions assumes that groups of residues that form local interacting communities are correlated and switch their conformational states cooperatively.

**Figure 9.**
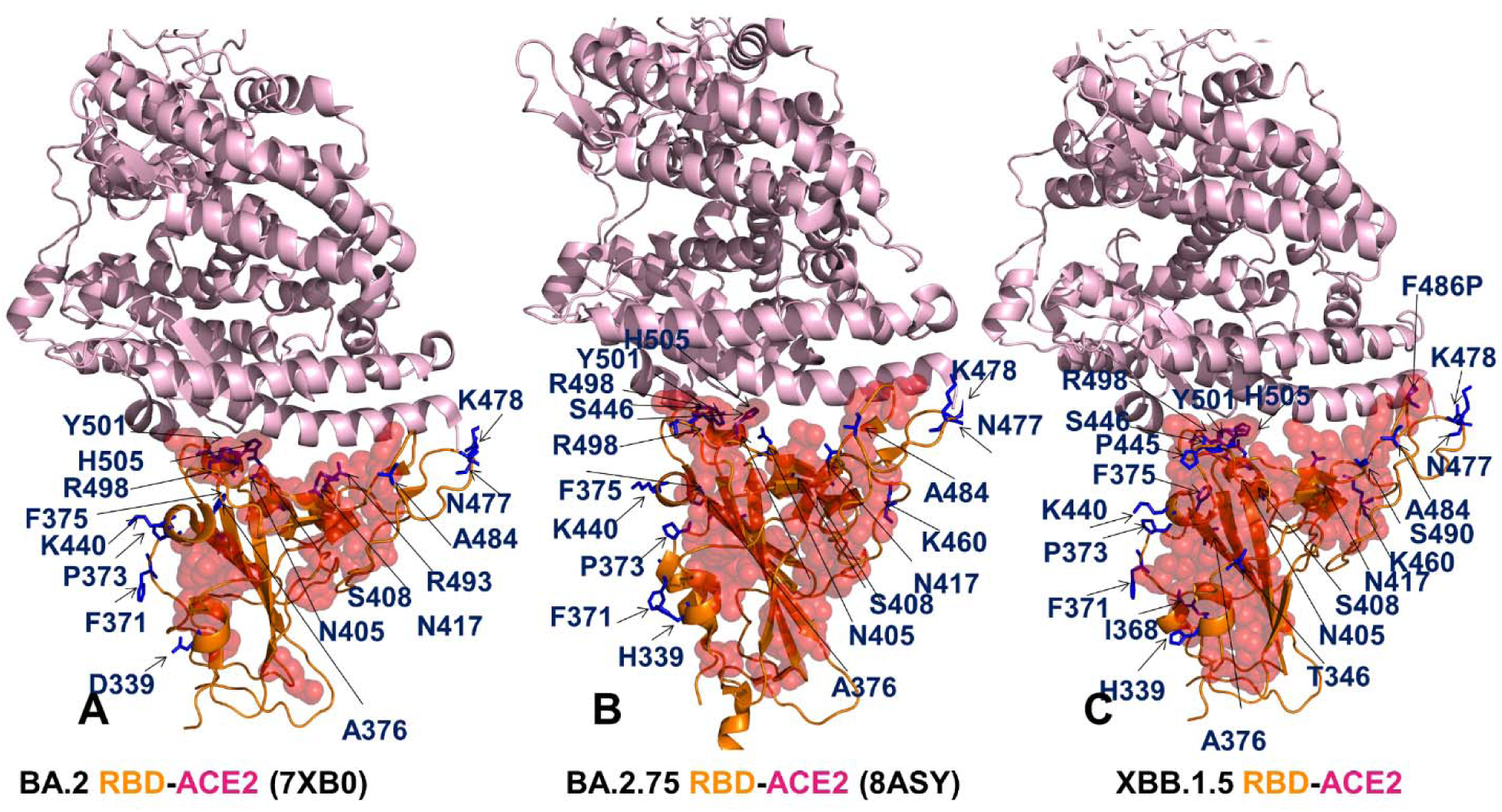
Structural mapping of the intrinsic RBD communities for the Omicron RBD BA.2-ACE2 complex, pdb id 7XB0 (A), the Omicron RBD BA.2.75-ACE2 complex, pdb id 8ASY (B), and the Omicron RBD XBB.1.5-ACE2 complex (C). The RBD is shown in orange-colored ribbons., ACE2 is in light, pink-colored ribbons. The RBD communities are shown in red-colored spheres with a 50% reduced transparency. The Omicron BA.2, BA.2.75 and XBB.1.5 mutational sites are shown in blue-colored sticks, annotated and indicated by arrows.

We first examined the distribution of the RBD communities in the RBD-ACE2 complexes assuming that the strength and the number of the intrinsically stable RBD modules provides an estimate of the folding stability (Figure 9). Structural mapping of the RBD communities revealed an appreciable and progressive increase in the RBD communities from the BA.2-ACE2 complex (Figure 9A) to BA.2.75 (Figure 9B) and XBB.1.5 (Figure 9C). The analysis showed a broadly distributed and dense network of inter-connected stable communities in BA.2.75 and XBB.1.5 complexes featuring a number of commonly shared modules. Of particular interest was to examine the involvement of the Omicron mutational sites in the RBD communities which could shed some light on their role in mediating the RBD stability. Although the key binding role of R498, Y501 and H505 sites is well-established, their role in mediating intrinsic RBD stability is less obvious. Strikingly, we found that in the BA.2 complex none of the Omicron sites contributes to the intrinsic RBD communities, and structural mapping illustrated the peripheral location of these sites relative to the RBD communities (Figure 9A). At the same time, in the XBB.1.5 complex, N417 and I368 mutational sites are involved in large and stable RBD clusters (Figure 9C). N417 is linked in one community with Y453-V350-W353-I402-I418-N422-Y423-Y495-D398-F400 which includes some of the most stable RBD positions F400, I402, I418 and Y453. This community becomes fragmented in less stable XBB.1 variant. In addition, while L368 is involved in local community F338-F342-L368 in BA.2, the mutational site I368 helps to consolidate stable clusters in the XBB.1.5 RBD-ACE2 complex that include a group of hydrophobic RBD core residues I358, V524, F338, F342, Y365, I368, V395, V513, F515, I434, F392, F374 and F377 residues (Figure 9C). In general, the results emphasized that Omicron mutations play a relatively minor role in mediating RBD stability, showing a significant tolerance to substitutions without deleterious impact on the RBD folding stability. The central finding of this network-based community analysis is a clear trend showing the increased density and spatial “expansion” of the stable RBD communities in the BA.2.75 and especially XBB.1.5 complexes (Figure 9). Hence, we argue that these results support the notion that the improved viral fitness of the XBB.1.5 variant may arise from the enhanced RBD stability and stronger ACE2 binding while maintaining the favorable immune escaping profile by utilizing functional benefits of conformationally adaptive Omicron sites.

We further proceeded by examining and mapping the RBD-ACE2 interfacial communities (Figure 10). Due to structural and topological similarity of the binding interfaces, all RBD-ACE2 complexes share a number of major common communities that persist throughout simulations and drive binding thermodynamics. These interfacial communities are present in the BA.2 RBD-ACE2 complex and include D38-Y49-R498, Q24-Y83-N487, Y489-F456-K31, N330-D355-T500, D355-T500-Y41, R493-H34-Y453, Y41-K353-Y501, K353-Y501-H505, Y41-Y501-R498 and Y489-K31-R493 (Figure 10A). Importantly, these communities are inter-connected and form the primary network of long-range couplings between interfacial residues. A close inspection of the interfacial communities in the BA.2.75 RBD-ACE2 complex showed the emergence of several additional stable modules Q24-G476-N487, Q24-Y83-F486, S19-Q24-N477 that give rise to stabilization of a larger multi-residue community Q24-Y83-G476-F486-N487 (Figure 10B). It is worth noticing that the strengthening of the binding interface network is mainly due to stabilization of local modules mediated by F486/N487 residues. In the XBB.1.5 complex, there is further strengthening and expansion of the key interfacial communities that remain stable throughout the entire course of simulations, including Y489-K31-F456, Q325-N439-Q506, N330-D355-T500-Y41-L45 and Y41-K352-Y501-H505-R498 (Figure 10C).

**Figure 10.**
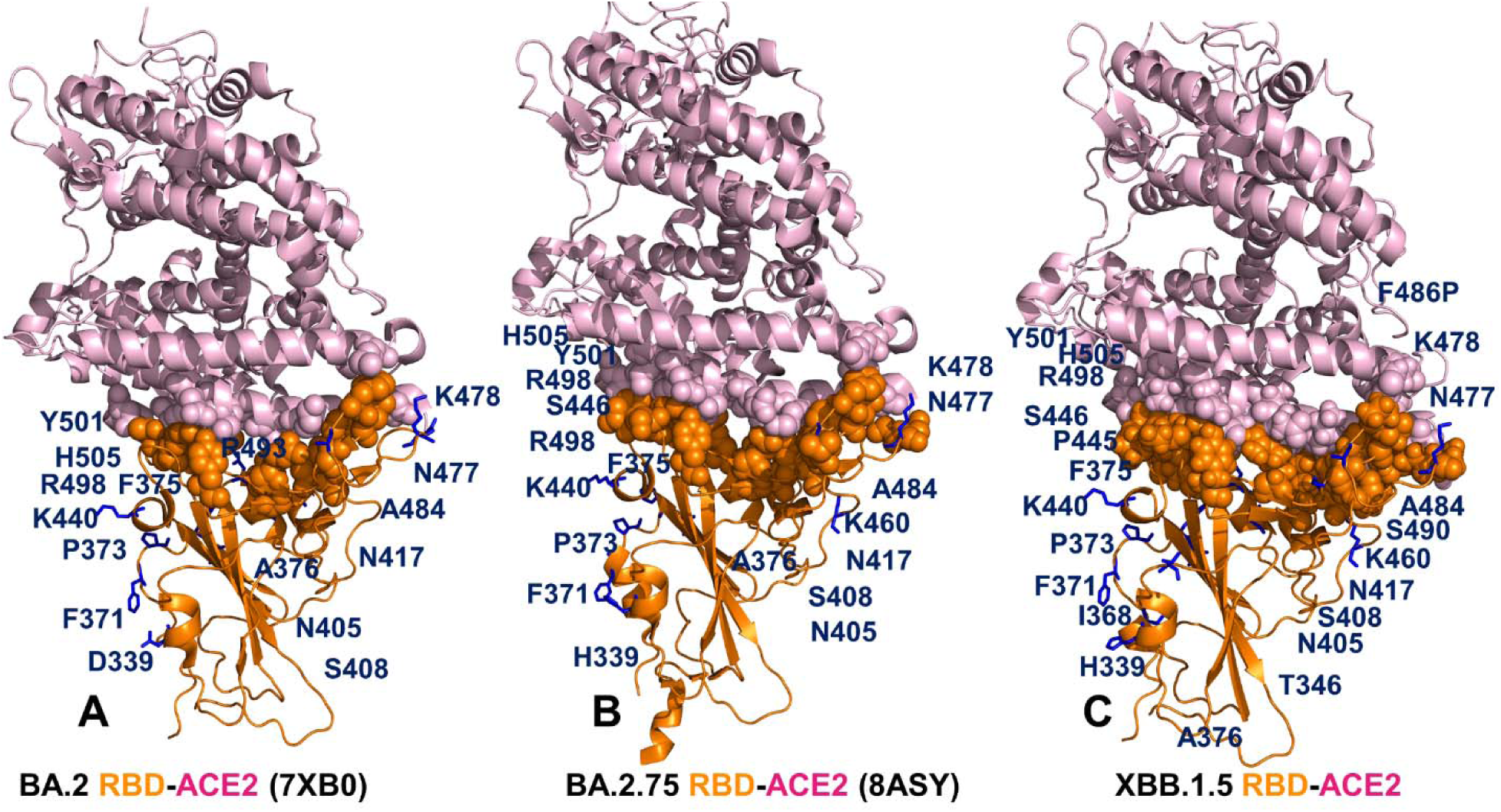
Structural mapping of the binding interface RBD-ACE2 communities for the Omicron RBD BA.2-ACE2 complex, pdb id 7XB0 (A), the Omicron RBD BA.2.75-ACE2 complex, pdb id 8ASY (B), and the Omicron RBD XBB.1.5-ACE2 complex (C). The RBD is shown in orange-colored ribbons., ACE2 is in light, pink-colored ribbons. The RBD-ACE2 communities are shown in spheres (RBD residues in orange and ACE2 residues in pink). The Omicron BA.2, BA.2.75 and XBB.1.5 mutational sites are shown in blue-colored sticks, annotated and indicated by arrows.

Overall, the analysis pointed out gradual strengthening and the increased density of major interfacial communities in the BA.2.75 and XBB.1.5 variants as compared to BA.2 variant (Figure 10). These results are consistent with the experimentally observed stronger binding of BA.2.75 and XBB.1.5 variants while also suggesting that the evolution of binding affinities in these variants may be determined by long-range network interactions between the RBD interface residues. Importantly, the commonly shared RBD-ACE2 interfacial communities are primarily mediated by R498, Y501 and Y489 hotspots making them indispensable hubs of the global interfacial network and controlling long-range allosteric interactions across the binding interface. We argue that through these networks of local communities the R498 and Y501 binding hotspots may allosterically enhance binding strength in other “soft” mutational sites due to strong epistatic couplings which may allow for compensatory rebound of binding affinity losses caused by accumulation of immune evasion modifications [67].

To characterize and rationalize the experimentally observed epistatic effects of the Omicron mutations [65-67], we further explored the network analysis and proposed a simple clique-based network model for describing non-additive effects of the RBD residues for the ensemble-based distribution of stable and meta-stable residue interaction cliques. In network terms, a *k*-clique is defined a complete sub-graph of *k* nodes in which each pair of nodes is connected by an edge, reflecting strong mutual interactions and dynamic coupling between every node in the clique with all other nodes that belong to the same clique. A collection of all interconnected *k*-cliques in a given network defines a *k*-clique community. We assumed that if the mutational sites of the Omicron RBD complexes fall into to a 3-clique structure we can associate this topological organization as a signature of potential local non-additive effects. At the same time, if particular Omicron mutations induce global changes in the distribution and connectivity of 3-cliques along the RBD-ACE2 binding interface, these Omicron mutations may have long-range epistatic effects on other Omicron interfacial positions that can be propagated and communicated via allosterically coupled network of the interfacial 3-cliques. Using topological network analysis, we reported the overall distributions and composition of stable 3-cliques in the interfacial network that are induced by the key Omicron mutations by Q498R, N501 and Y505H known to be involved in strong compensatory epistasis (Figure 11). [67].

**Figure 11.**
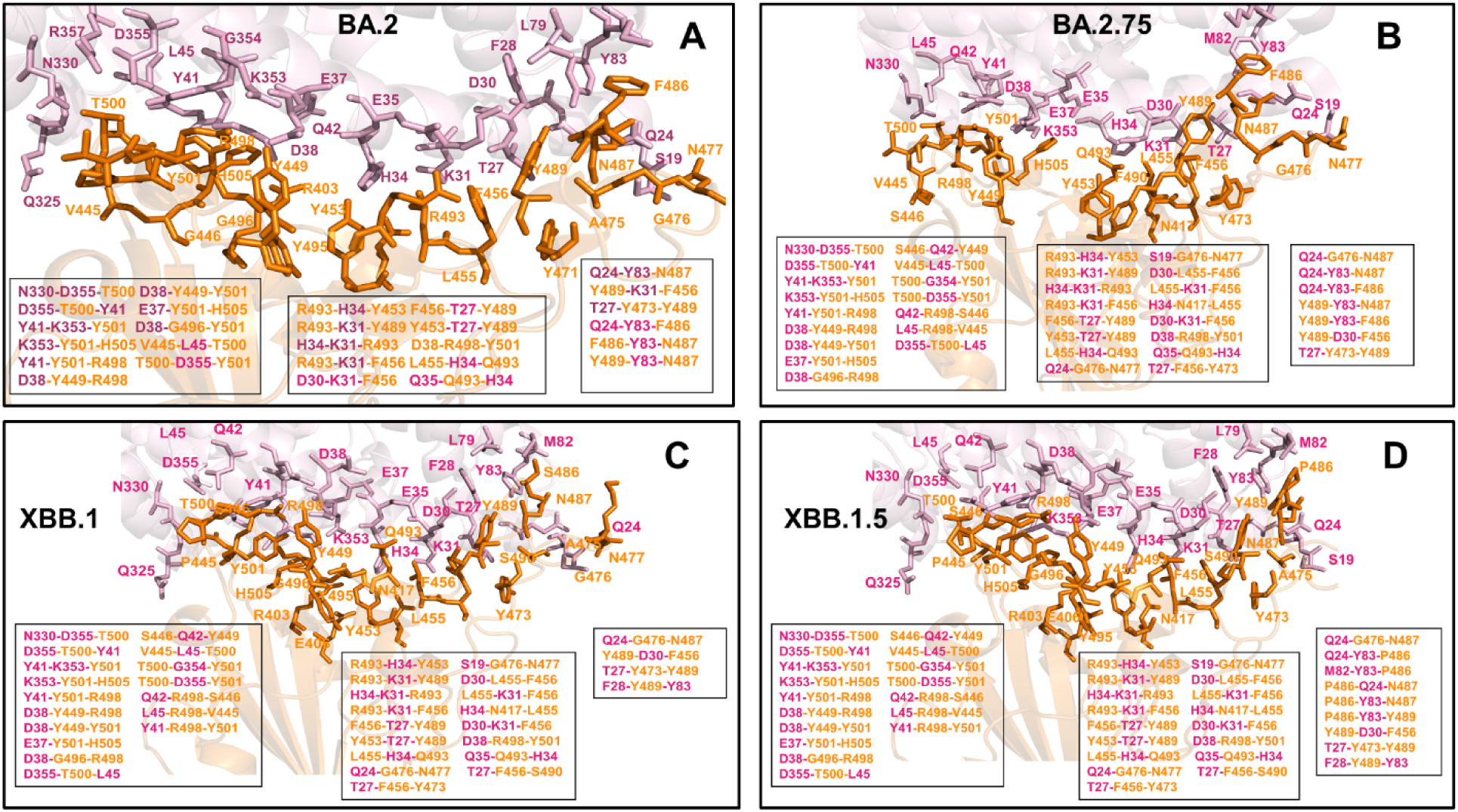
The network-based 3-clique analysis of potential epistatic relationships between RBD residues. The distributions of persistent 3-cliques formed at the binding interface of the Omicron RBD complexes with hACE2. Structural mapping and full annotation of the intermolecular 3-cliques for the RBD BA.2-hACE2 complex (A), the RBD BA.2.75-hACE2 complex (B), the RBD XBB.1-hACE2 complex (C) and the RBD XBB.1.5-hACE2 complex. The RBD binding interface residues are shown in orange sticks and the hACE2 binding residues are in cyan sticks.

Our results suggested that non-additive effects may depend on both structural topology and dynamic couplings between mutation sites, which can be quantitatively determined using the distribution of the inter-connected *3-*cliques. In the Omicron RBD BA.2 complex, we found a significant accumulation of stable interfacial 3-cliques most of which were directly anchored by Y501 and H505 mutations (K353-Y501-H505, D38-G396-Y501, Y41-K353-Y501, K353-Y501-H505, T500-D355-Y401 and Y41-Y501-R498) (Figure 11A). We also found that the interfacial 3-cliques anchored by Y501 are the most stable and persist throughout the course of simulations. In network terms, the involvement of N501Y in multiple 3-cliques implies that this mutational site not only enhances ACE2 binding but is strongly dynamically coupled with several RBD positions (Y449, G496, R498 and H505) and allows for strengthening of their binding interactions thus amplifying the effect of Omicron mutations. The network distribution also revealed that R498 and Y501 mutational sites can promote a larger number of stable 3-cliques at the central interfacial patch including D38-R498-Y501, R493-H34-Y453, R493-K31-Y489, H34-K31-R493, and R493-K31-F456 cliques (Figure 11A). Although most of these 3-cliques can be formed independently of N501Y and Q498R mutations, it appeared that an additional D38-R498-Y501 clique can be formed while the remaining cliques along the central patch become more persistent in simulations. These results highlighted the role of R493 position in anchoring multiple interaction clusters with the ACE2 residues, also indicating some level of dynamic coupling with Y453 and Y489 residues. The conserved Y453 and Y489 positions are involved in favorable hydrophobic interactions and dynamic coupling with the interactions mediated by R493 suggest some level of cooperativity and synchronicity between these contributions. At the same time, our analysis showed that the interfacial cliques in the distal flexible region of the interface (Q24-Y83-N487, Q24-Y83-F486, f486-Y83-N487, Y489-K31-F456, T27-Y473-Y489) are persistent independently of the presence of R498 and Y501 mutations (Figure 11A). Notably, we observed that F486 and N487 act as local mediators of stable cliques in this region.

Overall, while the topography of the RBD-ACE2 binding interface plays a fundamental role in determining the distribution and composition of the intermolecular network cliques, the dynamic residue couplings can modulate the strength and number of stable cliques through epistatic relationships. These observations agree with the functional studies showing that epistatic shifts in the RBD are primarily driven by Y501 site [65]. According to these experiments, the largest epistatic shift in mutational effects is associated with non-additive contribution of Q498R and N501Y, followed by less pronounced epistatic shifts at sites 491-496, 505-506, and 446-449 [65]. A considerably larger number of stable interfacial cliques are mediated by BA.2.75 Omicron mutations (Figure 11B), showing the increased number of cliques in the key binding region that are anchored by Y449, G496, R498, and Y501 mutations. Interestingly, we found that BA.2.75 variant may induce formation of additional stable cliques (Q42-R498-S446, L45-R498-V445, S446-Q42-Y449, V445-L45-T500) in which Y449 and R498 sites engage specific Omicron mutational positions V445 and S446 (Figure 11B). The denser network of inter-connected cliques in this region suggested that R498 and Y501 mutations may promote further strengthening of the binding interactions in the BA.2.75 variant amplifying the individually moderate effect of V445 and S446 mutations.

We also noticed the increased number of 3-cliques in a more dynamic region where several additional cliques are mediated by F486, N487 and Y489 positions thus suggesting a partial stabilization of the interfacial interactions in this region (Figure 11B). However, this consolidation of the binding interface in BA.2.75 occurred independently of the R498 and Y501 mutational sites and is mainly determined by reduced mobility in F486 and N487 positions. A similarly significant number of the interfacial cliques are mediated by Y449, G496, R498 and Y501 residues in the critical binding region for the XBB.1 (Figure 11C) and XBB.1.5 complexes (Figure 11D). Importantly, network modeling highlighted a noticeable drop in the stable 3-cliques formed in the flexible interface region for the XBB.1 variant, revealing that F486S mutation may not only compromise the strength of local binding interactions but also weaken the network of RBD interfacial contacts in this region (Figure 11C). Strikingly, the network of 3-cliques in this region is fully restored and further enhanced in the XBB.1.5 variant as F486P mutation can reduce the flexibility in the interfacial interactions, likely providing allosteric contribution to the improved binding affinity of XBB.1.5 (Figure 11D).

The findings of this analysis also suggested that the extent of non-additive epistatic contributions in the Omicron RBD BA.2.75 and XBB.1.5 complexes may be stronger than in the other subvariants (Figure 11). It is worth emphasizing that the stabilizing cliques in the critical binding region are anchored by the hotspots R498 and Y501 that also emerged as potential epistatic hubs that can mediate strong dynamic and energetic couplings with other RBD residues. We also found that Y449, G496, and T505 are dynamically strongly coupled with Y501 through a persistent network of the interfacial cliques, confirming that these residues could form a second group of important epistatic centers. These observations are in agreement with functional studies which showed that the major sites exhibiting epistatic shifts in the presence of Y501 include G446, Y449, G496, Y499, H505 residues [65,67]. Our results also indicated the presence of dynamic couplings between G446/Y449, R498/Y501 and R493 that may be important in mediating broad epistatic shifts which is consistent with the experimental data [67].

Hence, a clique-based network model can identify highly correlated and potentially non-additive mutational sites in the Omicron RBD complexes and distinguish them from other mutational sites that are less likely to experience epistatic shifts. These results provide a plausible rationale to the experimentally observed epistatic relationships in which mutations G446S, Q493R, and G496S individually reduce ACE2 binding but via strong epistasis with the pair R498/Y501 these losses can be fully compensated [67]. According to our findings, R498/Y501 can promote the formation of extensive stable network of inter-connected 3-cliques that enables to rescue weaker binding potential of other mutational sites by amplifying their binding contributions. The non-additive effects are ensured by chain of linked 3-cliques in which each pair of nodes/residues is connected by an edge, indicating a strong and intense mutual interaction among amino-acids on these nodes.

Another interesting finding of the network analysis is the role of F486, F486S and F486P mutations in BA.2.75, XBB.1 and XBB.1.5 variants respectively on modulating the density of the interfacial 3-cliques in the flexible interfacial region. Our analysis suggested that mutations of F486 can radically alter the strength and density of the interfacial cliques in this region and these F486-mediated changes are independent of the R498/Y501 mutations (Figure 11). We found that F486S can appreciably reduce both local and global binding interactions in this region, while F486P can remarkably restore binding not only via the improved local packing but also by promoting stabilization of 3-cliques formed by RBD residues P486 N487 and ACE2 residues Q24 and Y83 (Figure 11D). These findings are consistent with the experimentally established role of F486 as a critical evolutionary hotspot in which mutations at residue F486, such as F486V, F486I, F486S, have been recurring among prior Omicron subvariants.

To summarize, the network-based community analysis provided an additional insight to the mutational scanning data showing that mutational changes in these three positions may be coupled and lead to non-additive negative epistatic effects. However, importantly, these effects are secondary as the major non-additive contributions are likely to arise from presence of large number of stable cliques mediated by Y501 position. Our findings suggested that the extent of non-additive contributions to the binding affinity may be greater for the Omicron BA.2.75 and XBB.1.5 complexes that displayed the strongest binding affinity among the examined Omicron subvariants.

## Discussion

The results of this study provided molecular rationale and support to the experimental evidence that the acquisition of functionally balanced substitutions that optimize multiple fitness tradeoffs between immune evasion, high ACE2 affinity and sufficient conformational adaptability might be a common strategy of the virus evolution and serve as a primary driving force behind the emergence of new Omicron subvariants. Functional studies suggested that the evolutionary paths for significant improvements in the binding affinity of the Omicron RBD variants with hACE2 are relatively narrow and require balancing between various fitness tradeoffs of preserving RBD stability, maintaining binding to ACE2, and allowing for immune evasion [65-67]. These factors may limit the “ evolutionary opportunities” for the virus to adapt new mutations that markedly improve ACE2 binding affinity without compromising immune evasion and stability. As a result, it led to growing realization that evolutionary pressure invokes a complex interplay of thermodynamic factors to “designate” a privileged group of Omicron mutational hotspots that drive binding affinity with the ACE2, while allowing other Omicron sites to readily evolve immune escape capabilities with minor destabilizing liabilities. Moreover, some studies proposed that immune evasion may be a primary driver of Omicron evolution that sacrifices some ACE2 affinity enhancement substitutions to optimize immune-escaping mutations [36-39]. By examining forces driving the accelerated emergence of RBD mutations it was suggested that the immune pressure on the RBD becomes increasingly focused and promotes convergent evolution on the same sites including R346, K444, V445, G446, N450, L452, N460, F486, F490, R493, and S494 most of which are antibody-evasive [62]. These findings indicated that Omicron subvariants may evolve to accumulate convergent escape mutations while protecting and maintaining mutations that enable sufficient ACE2-binding capability. Analysis of the convergent evolution provided a useful summary of the observed tuning of the ACE2 binding affinity seen in new Omicron sub-lineages [63]. In particular, XBB.1 features E484A inherited from the BA.2 parent but further mutated into A484T in the child XBB.1.3 while XBB.1.5 adopted F486P mutation (F486S in XBB.1), and BA.2.75.2 inherited F486S from BA.2.75 but further mutated into F486L in the child CA.4 [63]. Remarkably, convergent evolution seen in these examples allowed for the improved or neutral binding affinity changes and immune escape. Several lines of evidence indicated that the observed coordination of evolution at different sites is largely due to epistatic, rather than random selection of mutations [140]. Epistatic interactions between mutations add substantial complexity of their adaptive landscapes are believed to play an important role in the virus evolution. The proposed in our study a network-based model for the analysis of non-additive contributions of the RBD residues indicated that some convergent Omicron mutations such as G446S (BA.2.75, BA.2.75.2, XBB), F486V (BA.4, BA.5, BQ.1, BQ.1.1), F486S, F490S (XBB.1), F486P (XBB.1.5) can display epistatic relationships with the major stability and binding affinity hotspots which may allow for the observed broad antibody resistance induced by these mutations [68]. Using atomistic simulations, the ensemble-based mutational scanning of binding/stability and network-based approaches, we showed that the binding affinity hotspots R498 and Y501 serve as central mediators of the interfacial communities in the RBD-ACE2 complexes. As a result, epistatic couplings mediated by R498 and Y501 hotspots with the RBD residues 491-496, 446-449 can be exemplified at the network level as these positions are involved in strong independent interactions within stable network cliques. This may allow for moderate negative effects on ACE2 binding in various Omicron immune evasion sites to be mitigated by strong compensatory epistasis exhibited by Y501. An important lesson of this study is the spatially localized dependence of mutational effects on preexisting mutations R498 and Y501. According to the experimental data, while the effect of V445P and G446S mutations in XBB.1/XBB.1.5 can be compensated through epistatic couplings with R498/Y501, a single mutation F486S in XBB.1 results in an appreciable loss of binding affinity and could not be offset by presence of the background R498/Y501 pair. Our results suggested that the restored binding strength mediated by F486P mutation in XBB.1.5 variant may arise from a spatially localized redistribution of local network cliques in the flexible interface region which is located on the other side of the binding interface from Y501 position. In global epistasis, the fitness effect of a particular mutation can be determined by the fitness of its genetic background. According to presented evidence, pairs of Omicron substitutions with the strong epistasis tend to be spatially proximal and form localized stable modules allowing for compensatory energetic changes. Previous studies noted similar patterns of epistatic substitutions co-occurring spatially more frequently than expected if the substitutions had occurred randomly [141]. Statistical sequence-based landscape analysis based on the direct coupling analysis (DCA) incorporated pairwise epistatic terms which allowed to capture local evolutionary constraints specific to the SARS-CoV-2 sequence background and identify K417, N440, E484, Q493, Q498, and N501 as sites of mutational enrichment [142]. Other studies have also found longer-range signals of co-occurring substitutions where allostery and epistasis conspire to facilitate evolution of new functions through coordinated mutations at distal sites [143]. The results of this study are consistent with the idea that protein stability in key sites can promote evolvability via epistasis and enhance tolerance to destabilizing mutations that often contribute to immune escape [137-139]. Overall, the interactions between Omicron mutational sites may be controlled by spatially localized compensatory epistatic relationships in which key binding hotspots can rescue binding affinities to offset the effects of destabilizing immune escape mutations.

## Conclusions

In this study, we systematically examined conformational dynamics, stability and binding of the Omicron RBD BA.2, BA.2.75, XBB.1 and XBB.1.5 RBD complexes with ACE2 using multiscale molecular simulations, in silico mutational scanning of the RBD residues and network-based community analysis of allosteric communications and epistatic interactions. Using multiscale simulation approach and conformational landscapes derived from all-atom MD simulations, we found a progressive rigidification of the RBD-ACE2 binding interface regions in the BA.2.75 and XBB.1.5 variants as well as the improved stability of the RBD core regions. The results of simulations showed that the improved RBD stability in BA.2.75 and XBB.1.5 complexes with ACE2 may be one of the factors linked with the experimentally observed enhancement in their binding affinity with ACE2. By using distance fluctuations analysis of stability and allosteric propensities in the complexes, we found that the important RBD binding interface centers R498, Y501 and H505 featured high stability indexes, reflecting a considerable rigidification of these residues due to strong interactions with ACE2. Our results suggested that these RBD residues function as key stability centers, binding hotspots as well as allosteric mediators of long-range communications in all RBD-ACE2 complexes. A systematic mutational scanning of the RBD residues in the complexes with ACE2 identified a conserved group of protein stability centers and binding affinity hotspots that determine the binding thermodynamics. Our data provided support to the emerging mechanism that XBB lineage may have evolved to evade immune suppression and outcompete other Omicron subvariants through mutations of F486 which is a hotspot for establishing protective immunity against the virus. Consistent with the experimental data, our results revealed that functionally beneficial effects of destabilizing mutations may be contingent on compensatory effects provided by a binding hotspot cluster centered on the R498/Y501 positions. These findings highlighted the important feature of a mechanism in which key binding affinity hotspots R498 and Y501 enable compensatory epistatic interaction with other binding-neutral Omicron positions to balance tradeoffs between immune evasion, protein stability and ACE2 binding. To characterize and rationalize the experimentally observed epistatic effects of the Omicron mutations we explored the network analysis and proposed a clique-based network model for describing non-additive effects of the RBD residues. We found that the network analysis and community-based assessment of dynamic and energetic couplings can identify highly inter-dependent mutational sites in the Omicron RBD complexes that experience epistatic shifts. These results provided a plausible rationale to the experimentally observed epistatic relationships in which effects of Omicron mutations that can reduce ACE2 binding but are important for immune escape are compensated via strong epistatic interactions with the binding affinity hotspots R498 and Y501. The results of this study suggested distinct and yet complementary roles of the Omicron mutation sites forming a coordinated network of hotspots that enable efficient modulation and balance of multiple fitness tradeoffs including structural stability, host receptor binding, immune evasion and conformational adaptability which create a complex functional landscape of virus transmissibility.

## Author Contributions

Conceptualization, G.V.; methodology, G.V.; software, G.V., M.A. and G.G; validation, G.V.; formal analysis, G.V., M.A. and G.G.; investigation, G.V.; resources, G.V., M.A. and G.G.; data curation, G.V.; writing—original draft preparation, G.V.; writing— review and editing, G.V., M.A. and G.G.; visualization, G.V.; supervision, G.V.; project administration, G.V.; funding acquisition, G.V. All authors have read and agreed to the published version of the manuscript.

## Funding

This research was funded by KAY FAMILY FOUNDATION, grant number A20-0032.

## Data Availability Statement

Data is fully contained within the article. Crystal structures were obtained and downloaded from the Protein Data Bank (http://www.rcsb.org). All simulations were performed using NAMD 2.13 package that was obtained from website https://www.ks.uiuc.edu/Development/Download/. All simulations were performed using the all-atom additive CHARMM36 protein force field that can be obtained from http://mackerell.umaryland.edu/charmm_ff.shtml. The residue interaction network files were obtained for all structures using the Residue Interaction Network Generator (RING) program RING v2.0.1 freely available at http://old.protein.bio.unipd.it/ring/. The computations of network parameters were done using NAPS program available at https://bioinf.iiit.ac.in/NAPS/index.php and Cytoscape 3.8.2 environment available at https://cytoscape.org/download.html. The rendering of protein structures was done with interactive visualization program UCSF ChimeraX package (https://www.rbvi.ucsf.edu/chimerax/) and Pymol (https://pymol.org/2/) .

## Acknowledgments

The authors acknowledge support from Schmid College of Science and Technology at Chapman University for providing computing resources at the Keck Center for Science and Engineering.

## Conflicts of Interest

The authors declare no conflict of interest. The funders had no role in the design of the study; in the collection, analyses, or interpretation of data; in the writing of the manuscript; or in the decision to publish the results.

---

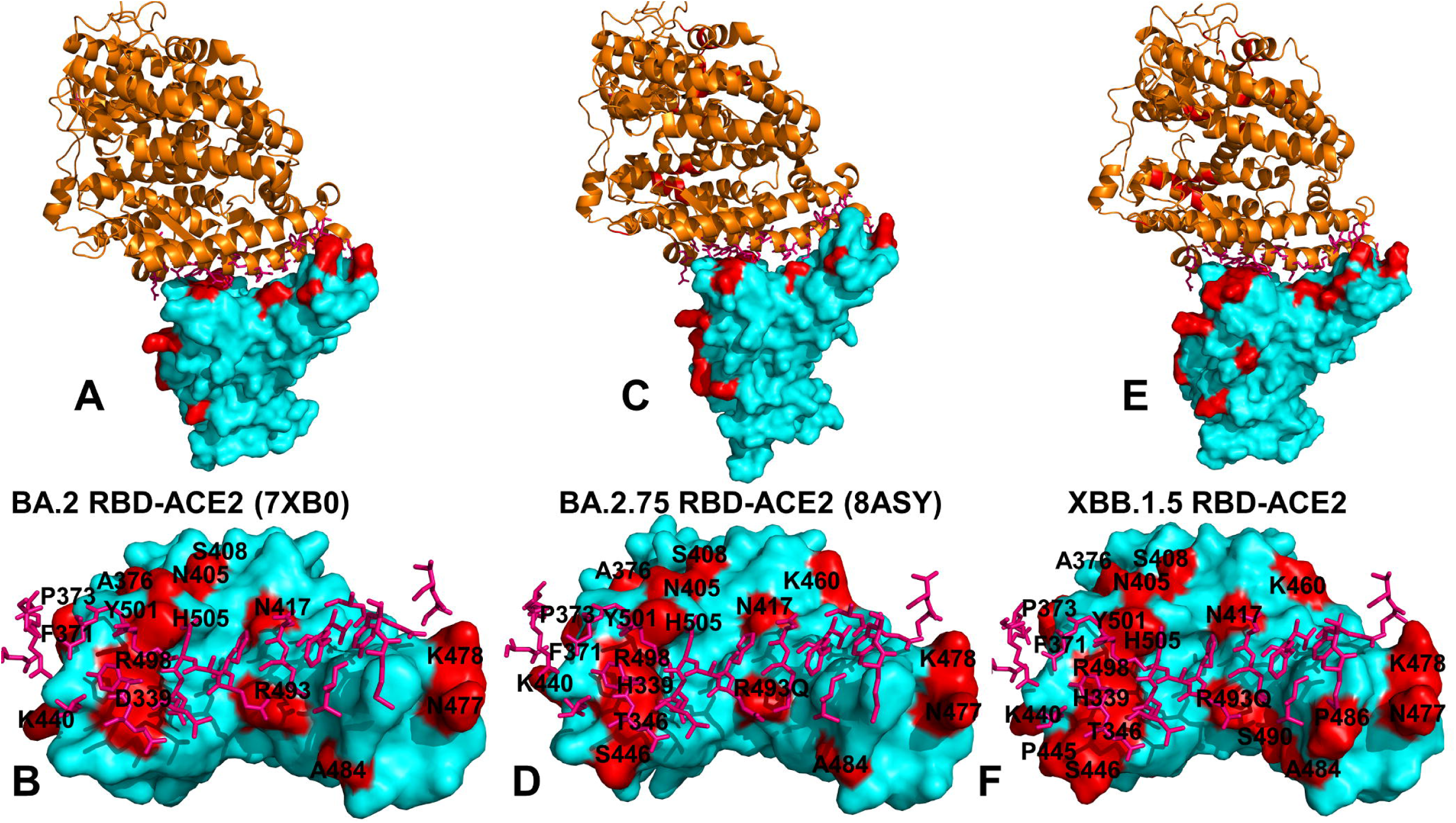

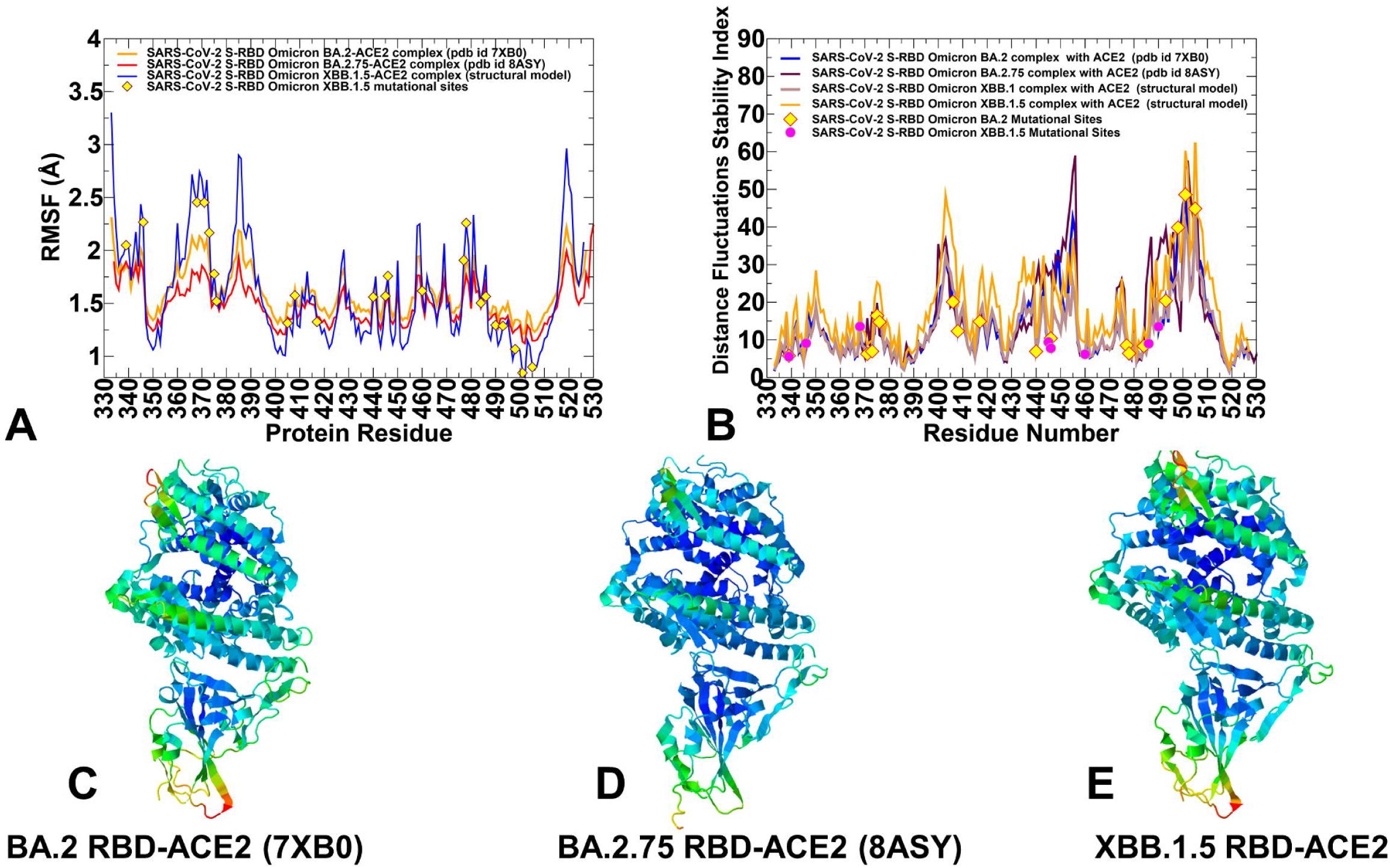

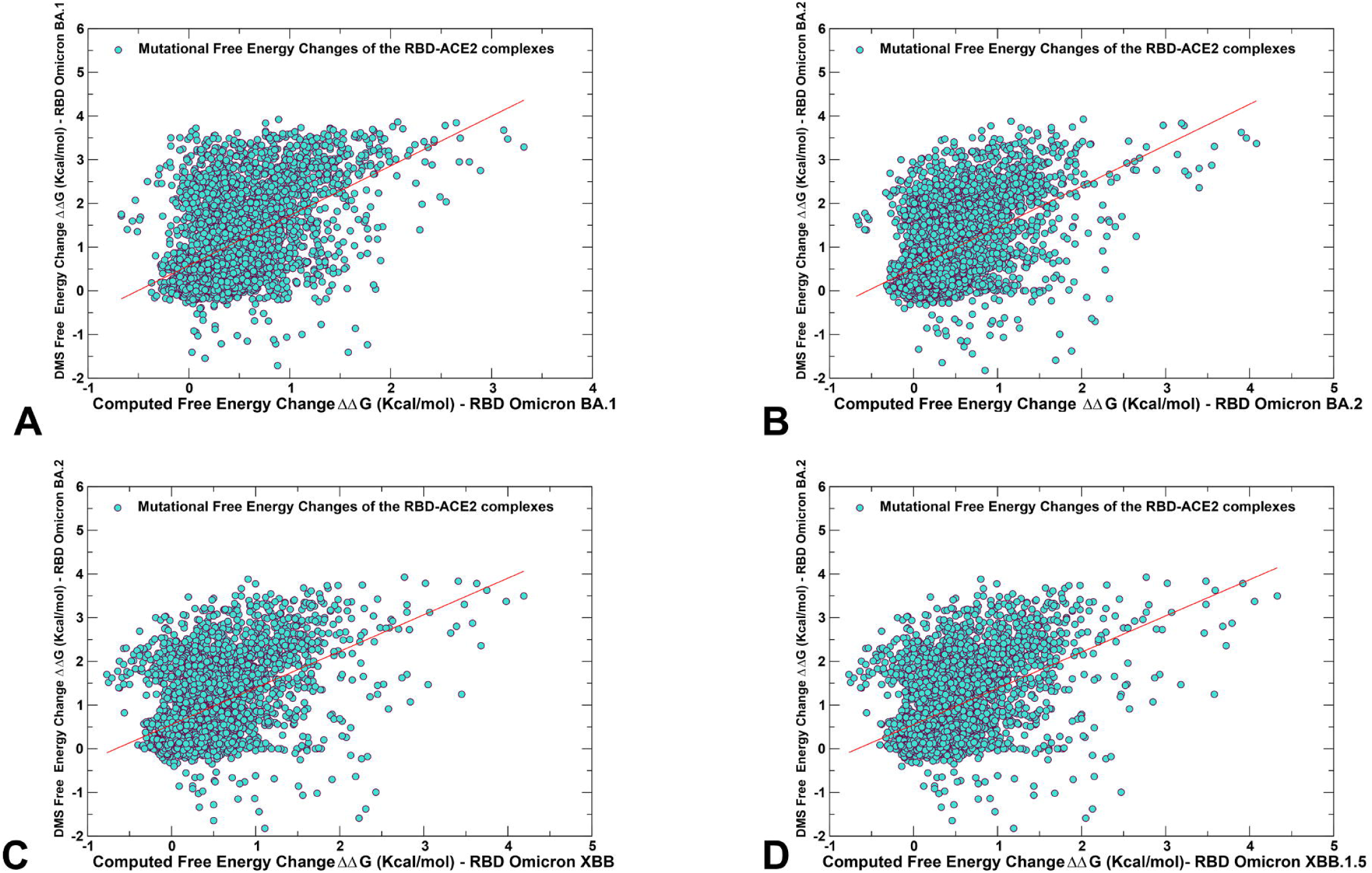

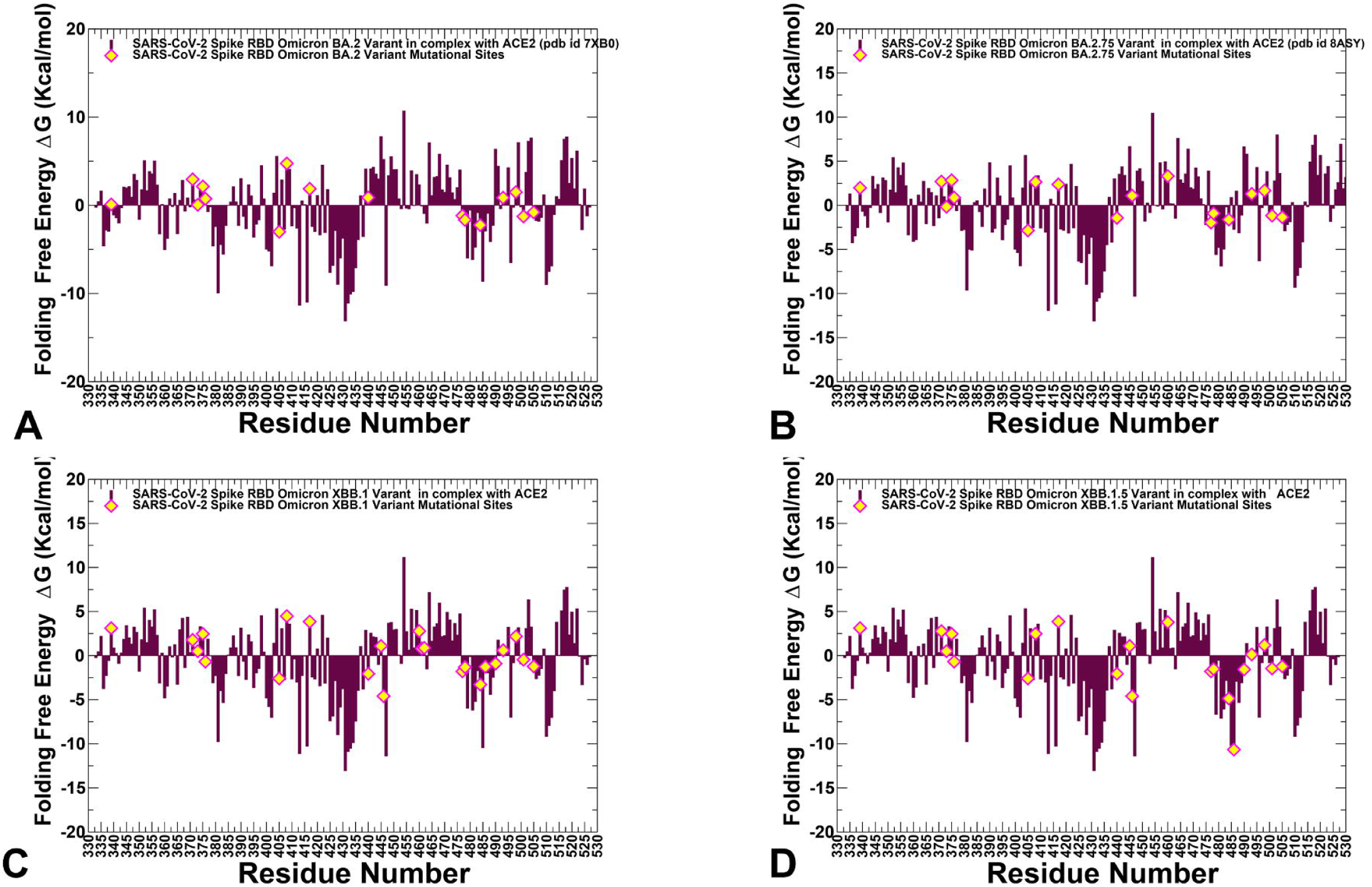

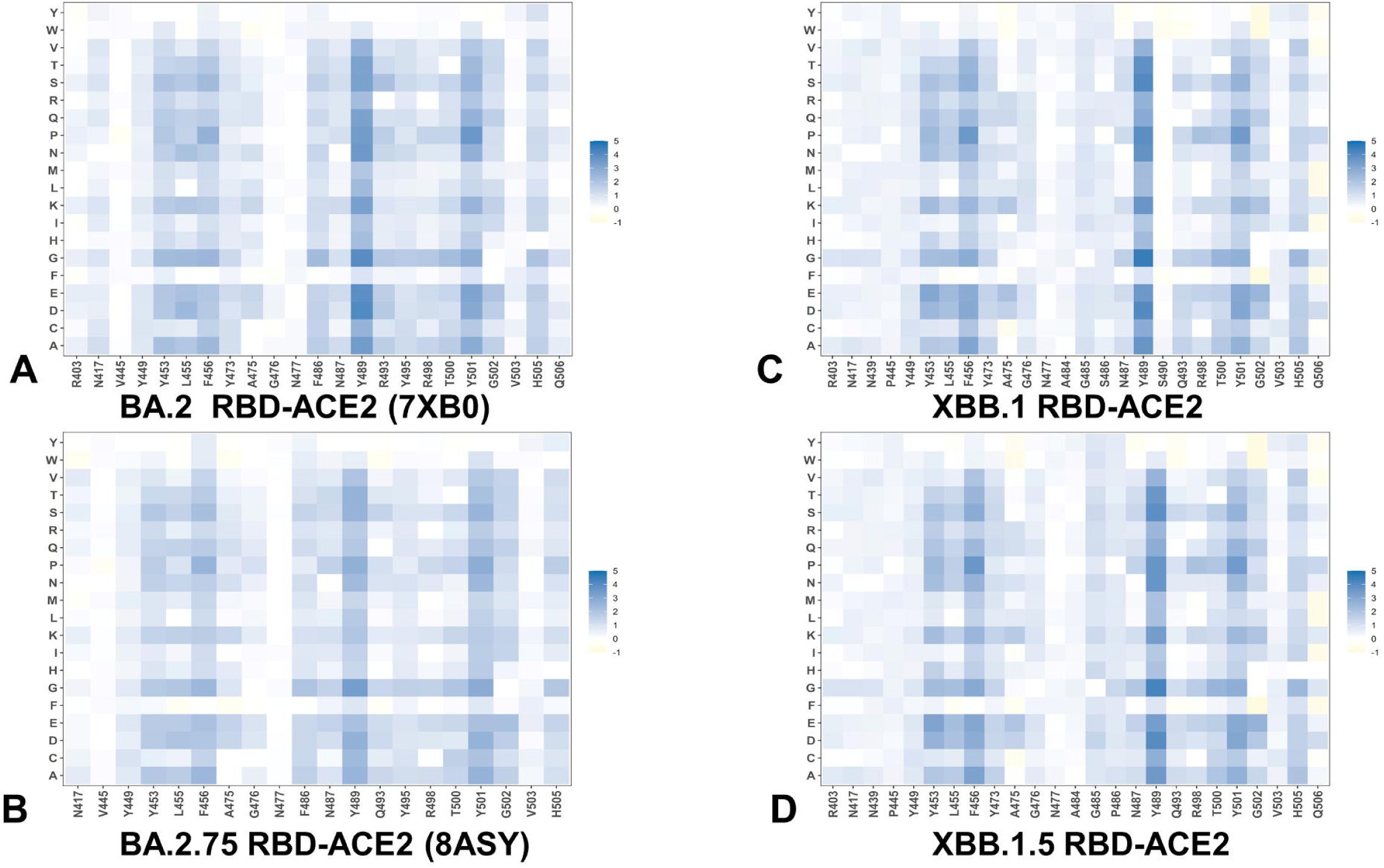

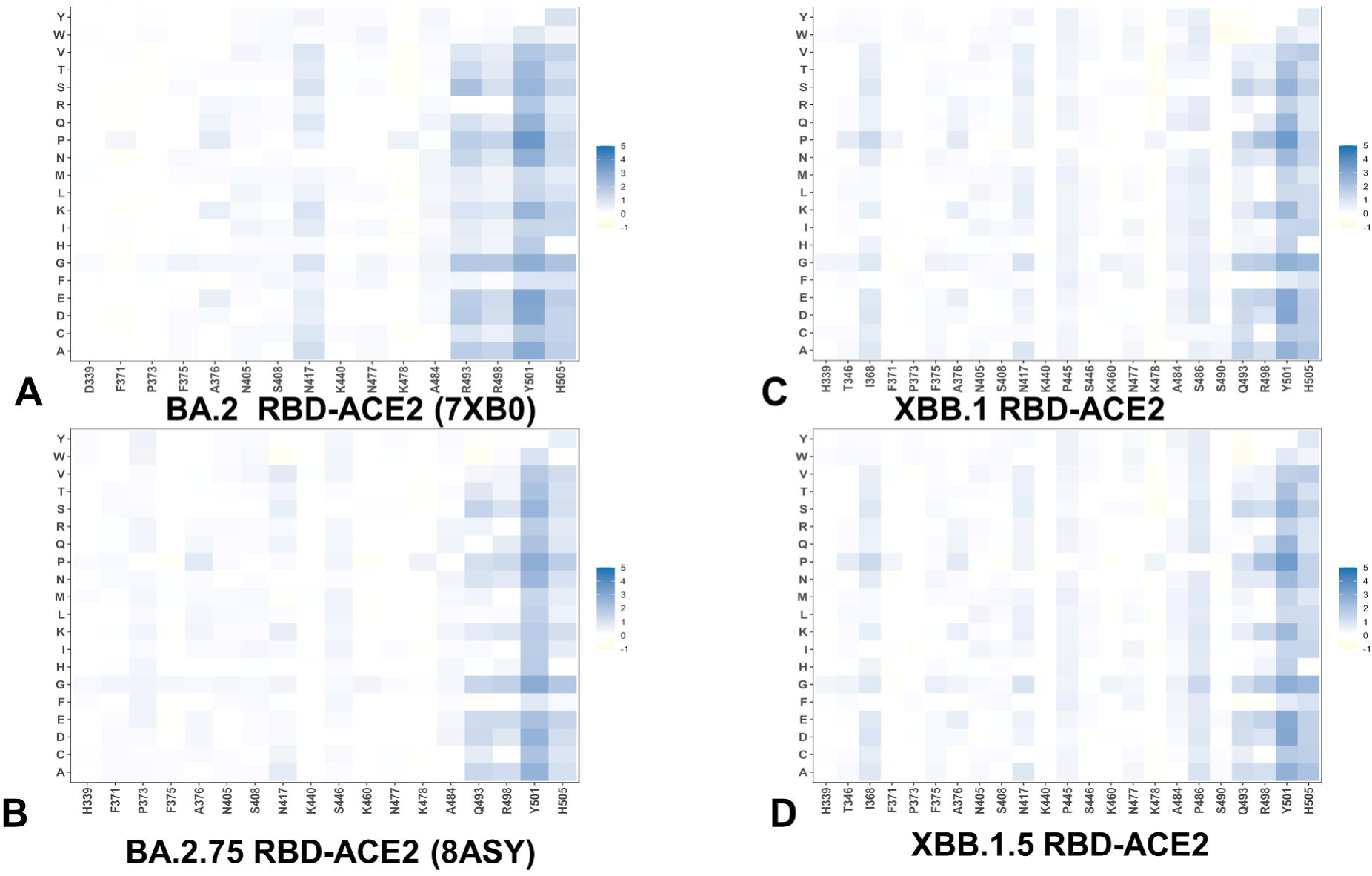

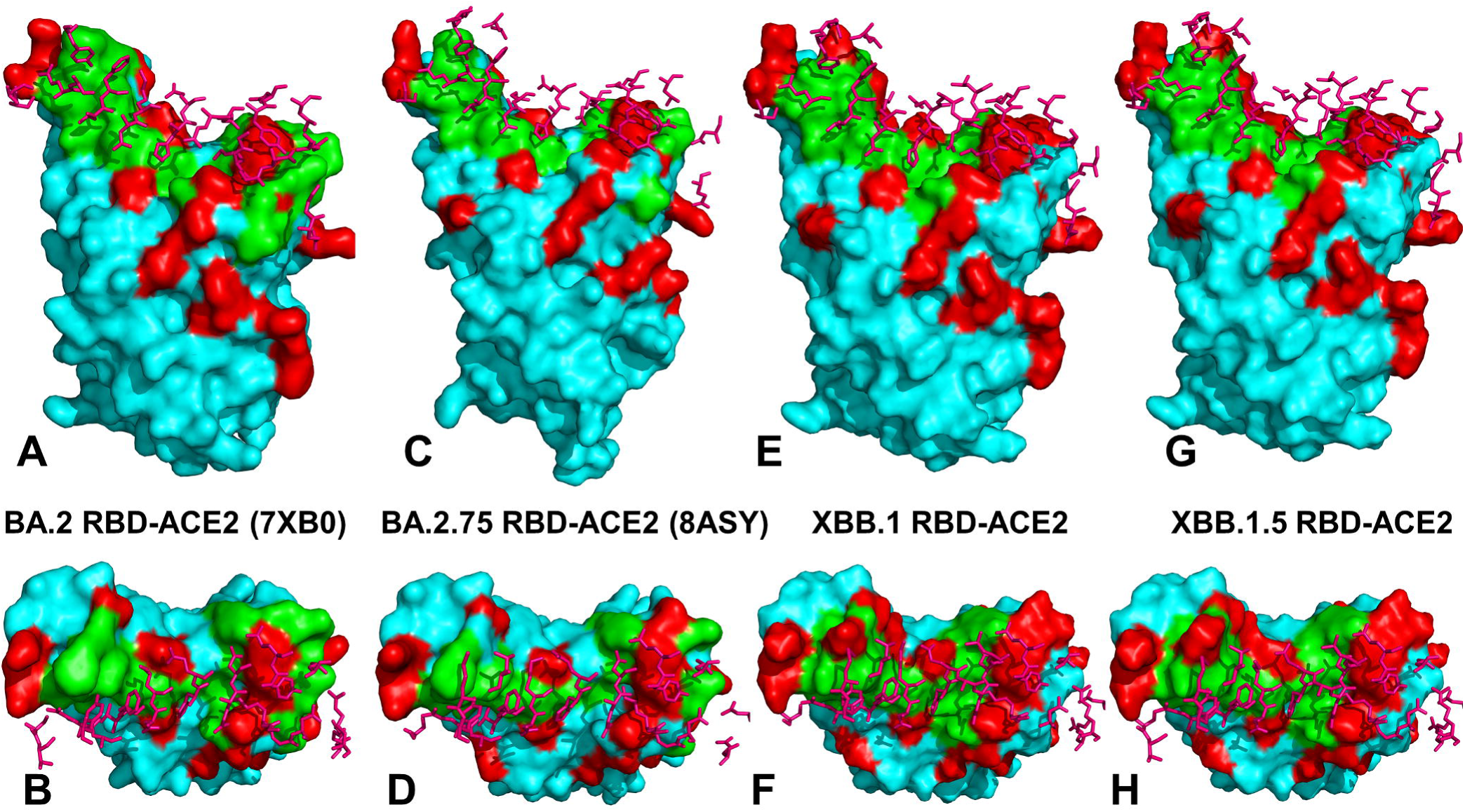

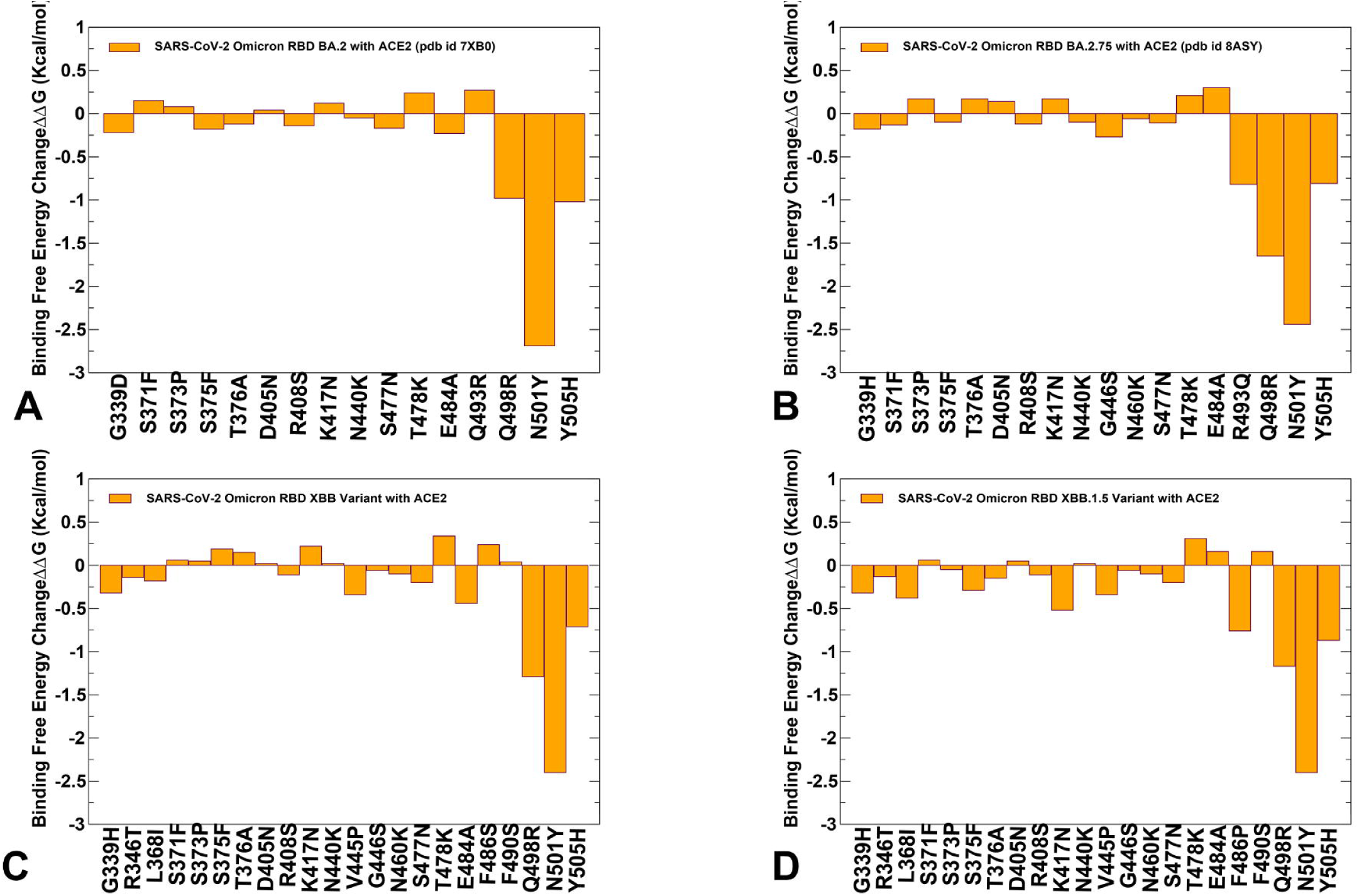

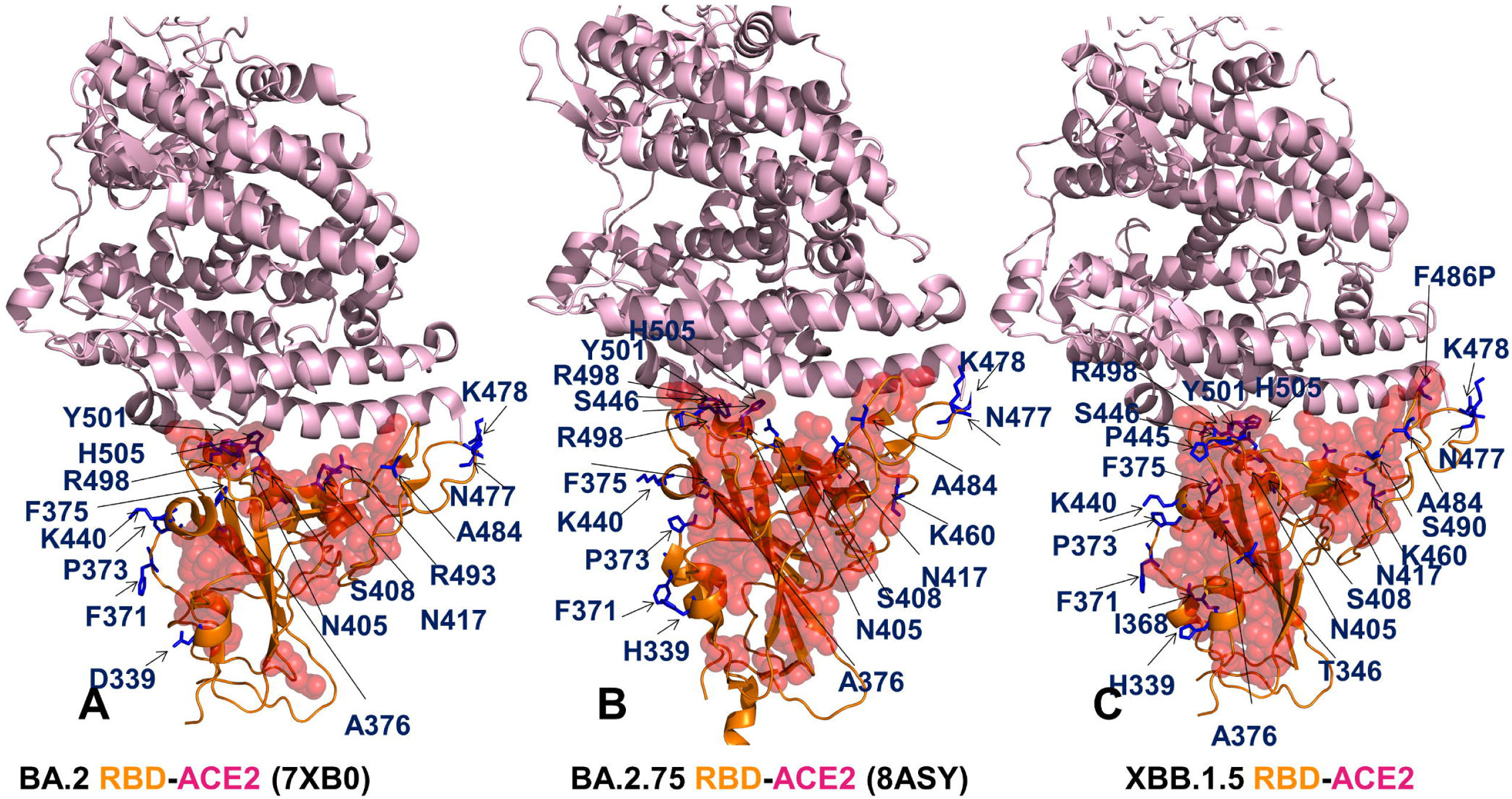

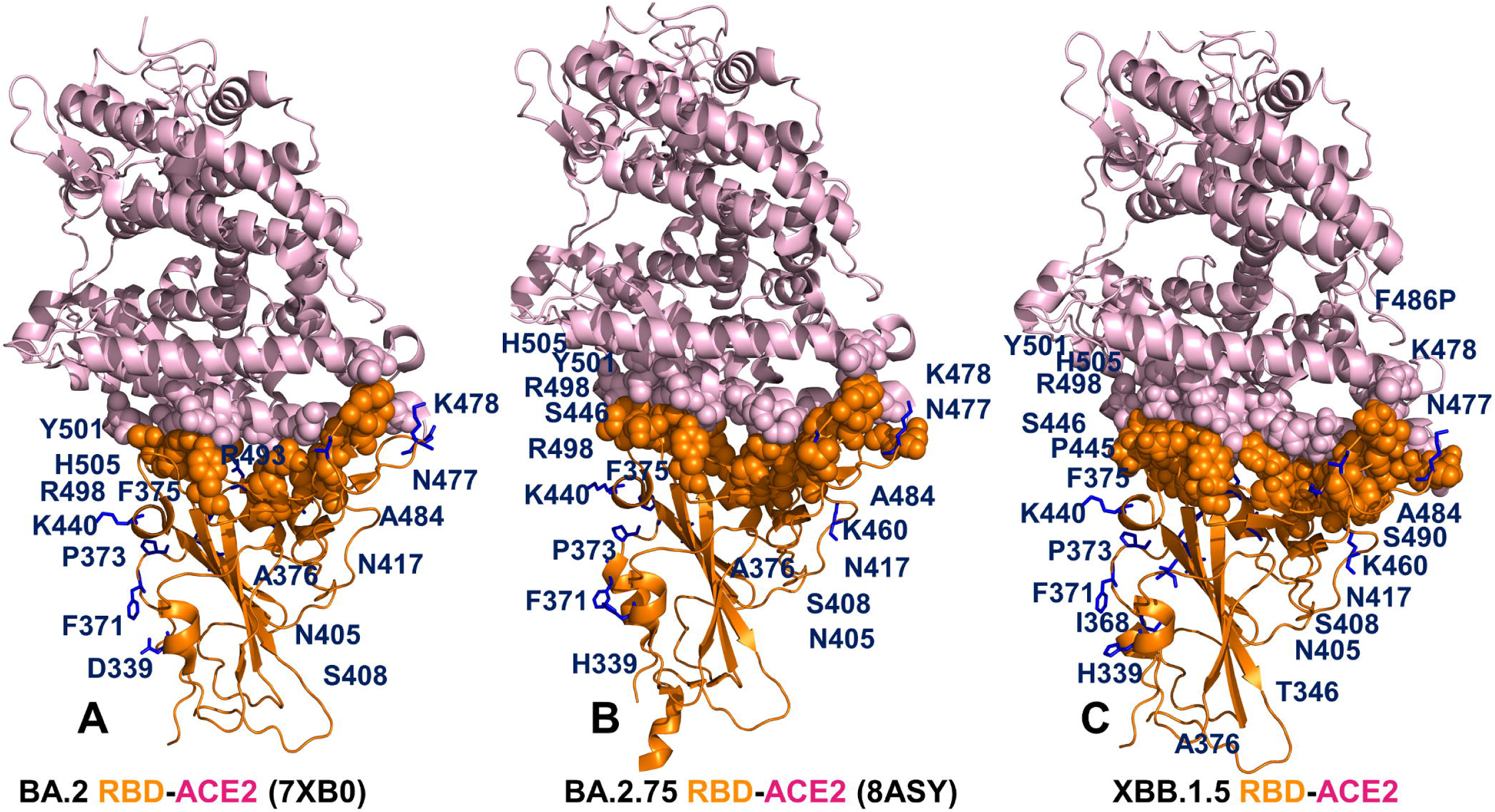

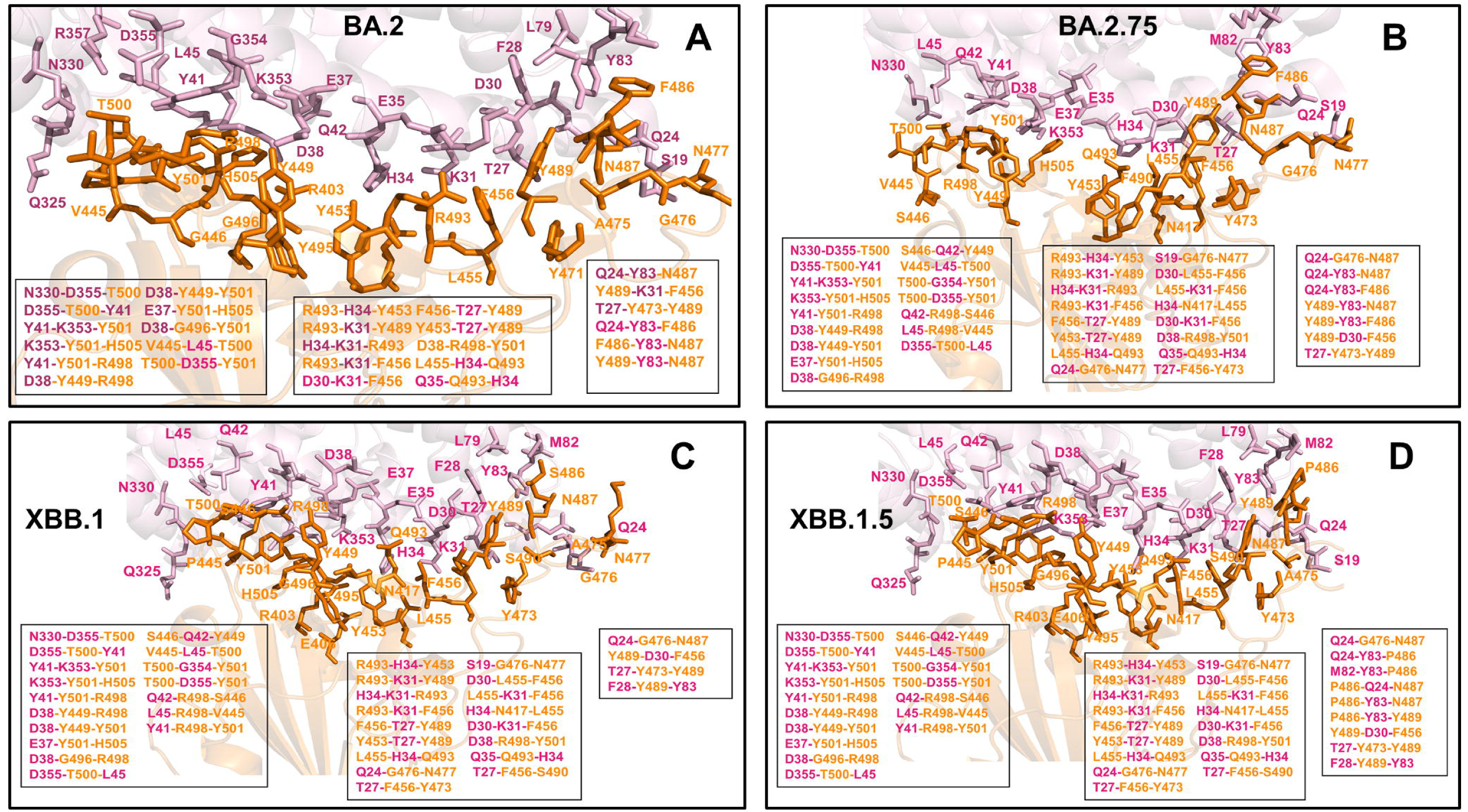

## Notes

### Competing Interest Statement

The authors have declared no competing interest.

